# Newly Evolved Endogenous Retroviruses Prime the Ovarian Reserve for Activation

**DOI:** 10.1101/2025.11.10.687753

**Authors:** Yasuhisa Munakata, Mengwen Hu, Raissa G Dani, Naokazu Inoue, Richard M. Schultz, Satoshi H. Namekawa

## Abstract

Female mammals are born with a finite pool of non-growing oocytes (NGOs) housed in primordial follicles, which form the ovarian reserve that determines reproductive lifespan. Mechanisms underlying the reserve’s long-term maintenance and subsequent follicular activation remain elusive. Using total RNA sequencing and de novo transcriptome assembly, we first captured the comprehensive oocyte transcriptome across perinatal oogenesis in mice. We show that NGOs establish accessible chromatin at gene regulatory elements—including promoters and enhancers—partly driven by newly evolved endogenous retroviruses (ERVs). In NGOs, epigenetic priming for follicular activation involves prior loading of transcription factors TCF3 and TCF12 and non-phosphorylated form of RNA polymerase II at these sites. This primed state is counteracted by repression via Polycomb Repressive Complex 1-mediated H2AK119 ubiquitylation. We propose that ERV-mediated epigenetic priming underlies the ovarian reserve’s long-term maintenance and establishes a transcriptionally competent yet repressive configuration that enables rapid gene activation upon oocyte growth.

## Introduction

Transposable elements (TEs) have coevolved with host genomes, which in turn exploit TEs to enhance reproductive fitness^1–3^. Many TEs function as gene regulatory elements—such as promoters and enhancers—that drive tissue- and cell type-specific gene expression^4,5^. The germline is a central arena in the evolutionary arms race between TEs and the host genome, where TE activity is finely tuned to support germline gene regulation^1,6,7^. Through this ongoing host–TE interplay, male and female germlines undergo distinct, unidirectional differentiation following sex determination, orchestrated by epigenetic priming^8,9^.

In the female germline, this differentiation culminates in formation of a finite pool of non-growing oocytes (NGOs) that reside within primordial follicles, whose maintenance sustains female fertility for decades in humans^10–13^. The size and quality of this follicle pool—known as the ovarian reserve—determine both fertility and reproductive lifespan. Prior to ovarian reserve formation, oocytes enter meiotic prophase I (MPI), during which homologous chromosomes synapse and recombine. Oocytes then arrest at the dictyate stage of MPI and form NGOs around birth. During the reproductive lifetime, the ovarian reserve progressively diminishes as primordial follicles are activated and enter the growth phase. Upon follicular activation, NGOs undergo massive transcriptional upregulation^14,15^ and transition to growing oocytes (GOs). GOs accumulate proteins, RNAs, and organelles that support early preimplantation development as they grow, ultimately becoming full-grown oocytes (FGOs) in preparation for maturation and fertilization^16–18^.

The transcriptome specific to GOs includes transcripts initiated from promoters of a subset of TEs, specifically long terminal repeat (LTR) retrotransposons^19–22^, many of which are not annotated as known genes^20^. This group comprises the autonomous Endogenous Retrovirus-L (ERVL) family and ERVL-apparent LTR retrotransposons (ERVL-MaLR)—a non-autonomous subclass originating from the ERVL family. Nevertheless, the full repertoire of oocyte transcripts and mechanisms that orchestrate genome-wide transcriptional activation—and the potential role of TEs in epigenetic priming—remain largely unexplored, largely due to the difficulty of isolating sufficient numbers of NGOs to comprehensively characterize transcriptomes and chromatin dynamics^8^.

To address this gap, we applied our recently developed pipeline for mouse NGOs^23–25^ to perform de novo transcriptome assembly and chromatin accessibility profiling across key stages of perinatal oogenesis—from MPI through ovarian reserve formation to oocyte growth. During ovarian reserve formation, NGOs acquire accessible chromatin at regulatory elements, including promoters and enhancers, many associated with newly evolved ERVs, specifically the ERVK family and ERVL-MaLR. We further identified an epigenetic priming mechanism in NGOs in which transcription factors TCF3 and TCF12^26^, together with RNA polymerase II, are already loaded to these sites but counteracted by Polycomb Repressive Complex 1–mediated H2AK119 ubiquitylation (H2AK119ub)^23,25^. These findings suggest that the ovarian reserve is actively maintained in a transcriptionally primed state and poised for later activation at the onset of oocyte growth.

## Results

### Identification of novel oocyte transcripts during perinatal oogenesis

In mice, ovarian reserve formation, which takes place around birth, is marked by several critical events that occur during a relatively short period of time, starting with germline cyst breakdown, followed by entry into dictyate arrest and assembly of primordial follicles (Fig. 1a)^27–30^. During this period, perinatal oocytes undergo the Perinatal Oocyte Transition (POT), during which they exit MPI and become NGOs^15,23^. A second major developmental transition, the Primordial-to-Primary Follicle Transition (PPT), occurs when a subset of NGOs is recruited to initiate oocyte growth. Both POT and PPT are characterized by extensive transcriptional reprogramming^15,31^.

**Figure 1.**
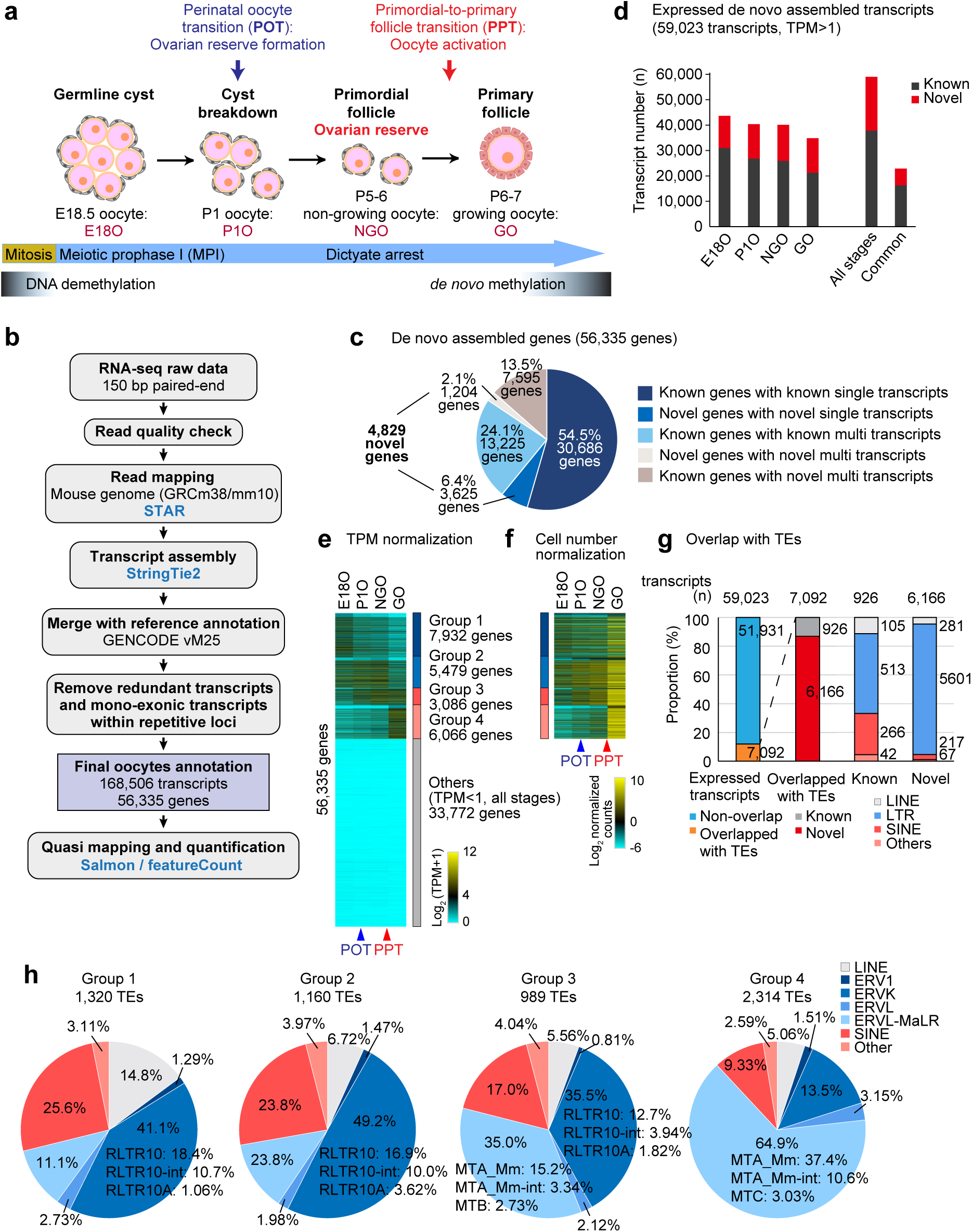
Assembly of the oocyte transcriptome during perinatal oogenesis. **a**, Schematic of mouse perinatal oogenesis and the four representative stages analyzed in this study: E18O, E18.5 fetal oocyte in germinal cyst; P1O, postnatal day 1 oocyte undergoing cyst breakdown during the perinatal oocyte transition (POT); NGO, non-growing oocyte in primordial follicles; GO, growing oocyte in primary follicles. DNA methylation levels are represented by color intensity. **b**, Overview of the strategy used for oocyte transcriptome assembly. **c**, Pie charts showing the composition of genes in the oocyte transcript annotation. **d**, Number of expressed de novo assembled transcripts at each stage. All stage: >1 TPM in at least one of the four developmental stages; Common: >1 TPM in all four developmental stages. **e**, Heatmap showing expression of expressed genes (>1 TPM in at least one stage) and low- or non-expressed genes (Others, <1 TPM in all stages) during perinatal oogenesis. Expressed genes were grouped into four k-means clusters. **f**, Heatmap showing gene expression normalized by ERCC spike-in across the four stages, based on the expressed genes identified in e. **g**, Bar charts showing the percentage of expressed genes overlapping with TSSs and TEs, and the percentage of TE families overlapping with TSSs of expressed genes. Overlaps were defined when a TE covered more than 51% of the 100 bp region surrounding the TSS. **h**, Pie charts showing the proportion of TE families overlapping with TSSs of expressed genes in each group defined in e.

To comprehensively define the full repertoire of oocyte transcripts and their gene expression dynamics during these major developmental transitions of perinatal oogenesis, we performed quantitative strand-specific total RNA sequencing (RNA-seq) and de novo transcriptome assembly at representative stages of mouse oogenesis. Specifically, we analyzed ribosomal RNA-depleted total RNA isolated from oocytes marked by the *Stella*-GFP transgene at four critical stages of development: embryonic day 18.5 (E18.5), corresponding to oocytes in the MPI stage residing in germline cysts (E18O); postnatal day 1 (P1), representing oocytes undergoing cyst breakdown during POT (P1O); P5 to 6, when oocytes have completed POT and reside in primordial follicles as NGOs; and P6 to 7, when oocytes have initiated growth and reside in primary follicles as GOs during PPT (Fig. 1a).

To enable per-cell resolution of transcriptome dynamics, we employed strand-specific total RNA-seq with ERCC spike-in controls^32^ to normalize expression to cell number. This approach addressed a major limitation of previous total RNA-seq studies in oogenesis, which lacked quantitative comparability across developmental stages due to the absence of normalization for input cell number^20,22^. The biological triplicates showed a high correlation within each developmental stage, ensuring reproducibility and robustness of the data (Extended Data Fig. 1a). After performing quality control and mapping the reads to the mouse genome (Fig. 1b), we assembled transcripts and classified the resulting de novo assembled transcripts into two categories: known transcripts, which overlapped with the GENCODE reference annotation, and novel transcripts, which did not (Extended Data Fig. 1b).

Our final oocyte transcript annotation consisted of 168,506 transcripts, including 28,210 mono-exonic and 140,296 multi-exonic transcripts, after exclusion of those originating from the Y chromosome and mitochondrial genome. In total, these transcripts were assigned to 56,335 genes (Fig. 1b, Extended Data Fig. 1b). In comparison to the 53,795 genes annotated in GENCODE (vM25, excluding Y-linked and mitochondrial genes), our assembly identified 4,829 novel genes (Fig. 1c). Notably, among all de novo-assembled genes, 13.5% (7,595 genes) were known genes that expressed previously unannotated multiple transcripts. Compared to a previous study that reported 82,939 oocyte transcripts from NGOs and GOs^20^, our dataset detected substantially more transcripts, as illustrated by representative genome browser views (Extended Data Fig. 1c).

To predict coding potential of novel transcripts, we used three independent programs: CPC2^33^, CPAT^34^, and CodAn^35^. The majority of novel transcripts were predicted to be non-coding (Extended Data Fig. 2a). Although the set of predicted coding transcripts varied across the three programs, 29 transcripts were consistently predicted to be coding by all three (Extended Data Fig. 2b). Many of these putative coding transcripts were located in intergenic regions (Extended Data Fig. 2c). Several showed partial homology to known proteins, including arf-GAP and HGH1 (Extended Data Fig. 2d, e). Although further characterization is needed, these results suggest that a subset of the novel genes may encode proteins homologous to known proteins.

Next, we defined 59,023 expressed transcripts as those with expression levels exceeding one transcript per million (TPM) in at least one of the four developmental stages, of which 21,150 were novel (Extended Data Fig. 3a). Remarkably, each developmental stage contained more than 10,000 novel transcripts (Fig. 1d). These novel genes exhibited robust expression, often at levels comparable to or exceeding those of known genes, particularly in GOs, although their expression was relatively lower in E18O, P1O, and NGOs (Extended Data Fig. 3b). These finding underscore the high abundance and dynamic regulation of novel transcripts throughout key stages of oogenesis.

Because multiple transcripts can arise from a single gene (e.g., through alternative promoter usage or alternative splicing), we next assessed gene-level expression to remove redundant read counts originating from the same locus, applying the same threshold (>1 TPM in at least one of the four developmental stages). Using this criterion, we identified 22,563 expressed genes, which were categorized into four groups based on k-means clustering of their expression patterns normalized by TPM (Fig. 1e), highlighting the two major transitions of POT and PPT. Group 1 genes were expressed during the MPI stage and enriched for GO terms associated with MPI processes (Extended Data Fig. 3c), while genes in Groups 2 and 3 were upregulated during POT and were associated with metabolic pathways and chromosome condensation, reflecting the transition from MPI to NGOs. To more accurately quantify expression dynamics per cell, which are often distorted by conventional normalization methods, we further normalized expression levels based on cell number using ERCC spike-ins, after excluding genes with very low or no expression (Fig. 1e, “Others”: 33,772 genes), as their signals were at background levels. This analysis revealed that the majority of detectable genes were upregulated in the PPT, quantitatively demonstrating the global transcriptional activation that follows follicle activation (Fig. 1f).

In GOs, a group of TEs—specifically certain families of ERVs—are highly transcribed and act as promoters for oocyte-specific transcripts^19–22^. Among the 59,023 expressed transcripts, 7,092 had transcription start sites (TSSs) overlapping with TEs in the same transcription direction; notably, 6,166 of these (86.9%) were novel transcripts (Fig. 1g). Of the TEs located at TSSs of novel transcripts, 86.2% were ERVs (Extended Data Fig. 3d). Within these, 42.9% were ERVKs (with RLTR10 comprising 19.8%) and 40.5% were ERVL-MaLRs (with mouse-specific MTA_Mm accounting for 22.1%) (Extended Data Fig. 3d). RLTR10 elements are newly evolved, mouse-specific ERVKs that emerged after the mouse–rat divergence approximately 12–14 million years ago^36^. MTA elements amplified in mice ∼2– 10 million years ago, younger than RLTR10s, likely expanding along the *Mus musculus* lineage^37^. A previous study identified 3,498 transcripts with TE-derived promoters during oogenesis^20^, and our findings corroborated and expanded this repertoire. Notably, TE usage varied across expression groups: ERVKs and SINEs were predominant in Group 1 and 2 genes, which were highly expressed in MPI and NGO stages, while ERVL-MaLRs dominated in Group 3 and 4 genes, which became active during the transition from NGOs to GOs (Fig. 1h). The changes reflect a progressive shift in promoter TE usage throughout oogenesis, particularly from MPI to GOs.

### The accessible chromatin landscape in perinatal oogenesis

To identify gene regulatory regions responsible for the dynamic shifts in gene expression that we observed, we performed omni-ATAC-seq^38^ (assay for transposase-accessible chromatin using sequencing) to identify accessible chromatin regions in perinatal oocytes during POT (Fig. 2a, Extended Data Fig. 4a). From E18O to NGO, we observed dramatic changes in chromatin accessibility (Fig. 2a), with a significant increase in the number of ATAC peaks, particularly within intronic and intergenic regions (Fig. 2b). The ATAC signal intensity remained largely unchanged in gene-proximal regions (±1 kb from TSSs), but progressively increased in gene-distal regions (beyond ±1 kb from TSSs) (Fig. 2c), where accessible chromatin is frequently bound by cell-specific transcription factors and functions as enhancers to regulate gene expression^39–41^.

**Figure 2.**
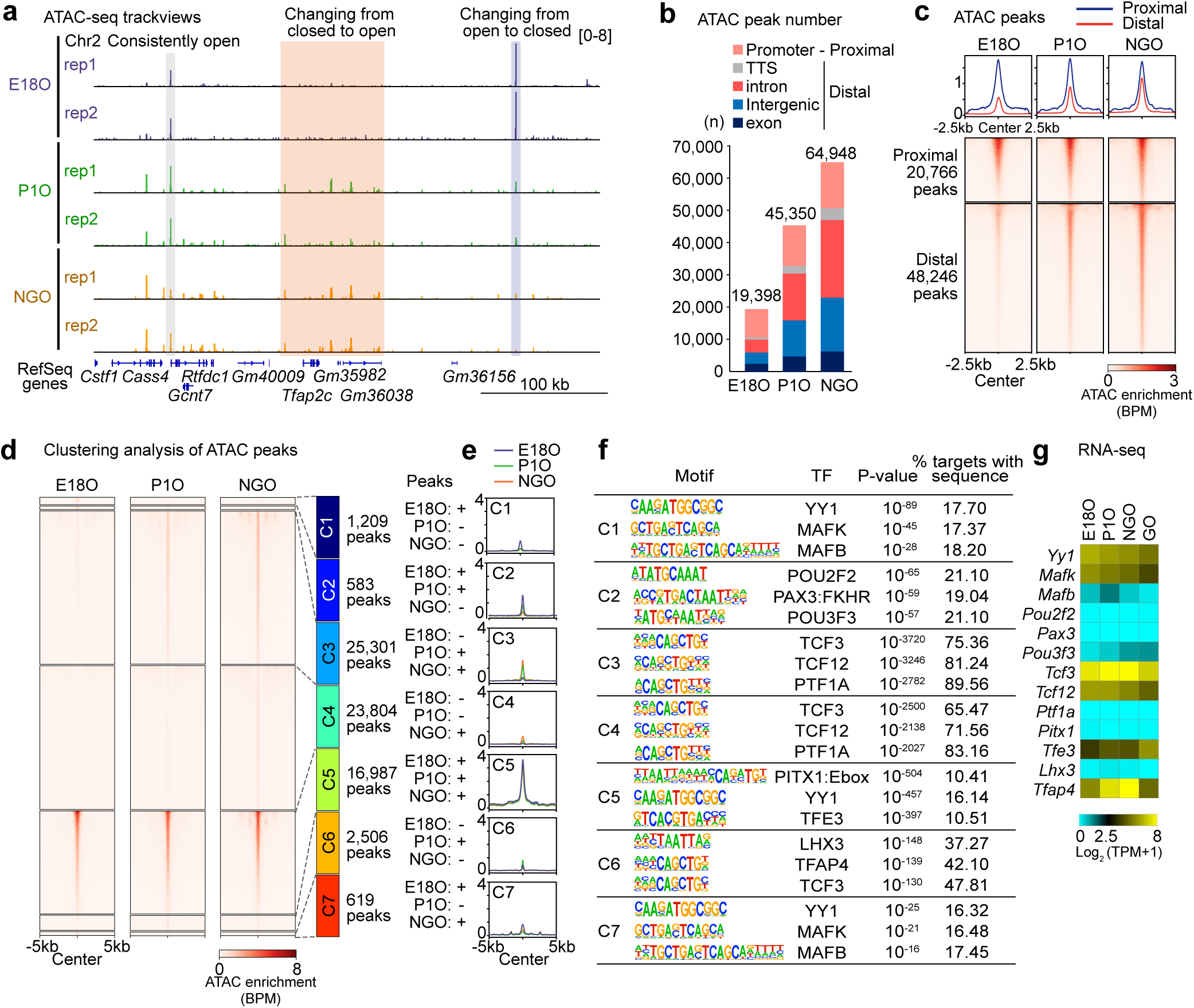
The landscape of accessible chromatin in perinatal oogenesis. **a**, Track views showing ATAC-seq enrichment in perinatal oocytes during POT. Data ranges are indicated in brackets. **b**, Numbers and genomic distribution of ATAC-seq peaks in perinatal oocytes during POT. Promoter: −1 kb to +100 bp from the transcription start site (TSS); TTS: −100 bp to +1 kb from the transcription termination site. **c**, Average tag density plots and heatmaps showing ATAC-seq enrichment dynamics at proximal (TSS ±1 kb) and distal (>1 kb from TSSs) peaks in perinatal oocytes during POT. **d**, Heatmap showing ATAC-seq enrichment dynamics for each cluster based on the presence of peaks in perinatal oocytes during POT. +: presence of peaks; –: absence of peaks. **e**, Average tag density plots showing ATAC-seq enrichment dynamics at peak regions (peak center ±5 kb) during perinatal oogenesis. **f**, HOMER known motif analyses of ATAC-seq peaks in each cluster defined in d. **g**, Heatmap showing RNA-seq expression levels of the transcription factors identified in f in perinatal oocytes during POT.

We further characterized the ATAC peaks through a clustering analysis, in which they were grouped into seven classes based on their enrichment profiles (Fig. 2d, e). Classes 3 and 4 showed increased accessibility, with Class 3 specific to P1O/NGO (25,301 peaks) and Class 4 to NGO (23,804 peaks), predominantly in distal regions from TSSs (Extended Data Fig. 4b). Motif analysis revealed enrichment of TCF3 and TCF12 binding sites (Fig. 2f) —basic helix–loop–helix (bHLH) transcription factors that regulate oocyte growth^26^—and both factors were consistently expressed from E18O to GO (Fig. 2g). In contrast, ATAC peaks in Class 5 (16,987 peaks) remained largely unchanged across developmental stages were primarily located in proximal regions near the TSSs (Extended Data Fig. 4b). These findings suggest that distal accessible regions, which might function as active regulatory elements, are established during POT.

In addition, proximal accessible regions exhibited changes correlated with gene expression. While proximal regions of expressed gene groups (identified in Fig. 1e) remained largely accessible throughout perinatal oogenesis, Group 1 genes—highly expressed in MPI—showed an overall reduction in ATAC signal intensity during POT (Fig. 3a), consistent with their decreased expression. Thus, although most TSSs remained constantly accessible, specific gene repression appeared to be associated with local chromatin closure. Notably, among the expressed genes, novel genes exhibited a gradual increase in accessibility at their TSSs and downstream regions from E18O to NGO, whereas TSS accessibility of known genes tended to decline progressively (Fig. 3b). A subset of novel genes displayed particularly high accessibility at downstream regions of TSSs (Cluster 1; 490 genes), 88.6% of TSSs in this cluster overlapped with TEs (Fig. 3c), and these downstream accessible regions were further enriched for TEs (Fig. 3d, Extended Data Fig. 4c). Collectively, these results suggest that a subset of novel genes, potentially driven by TEs, becomes specifically accessible during POT, preceding their robust expression in GOs.

**Figure 3.**
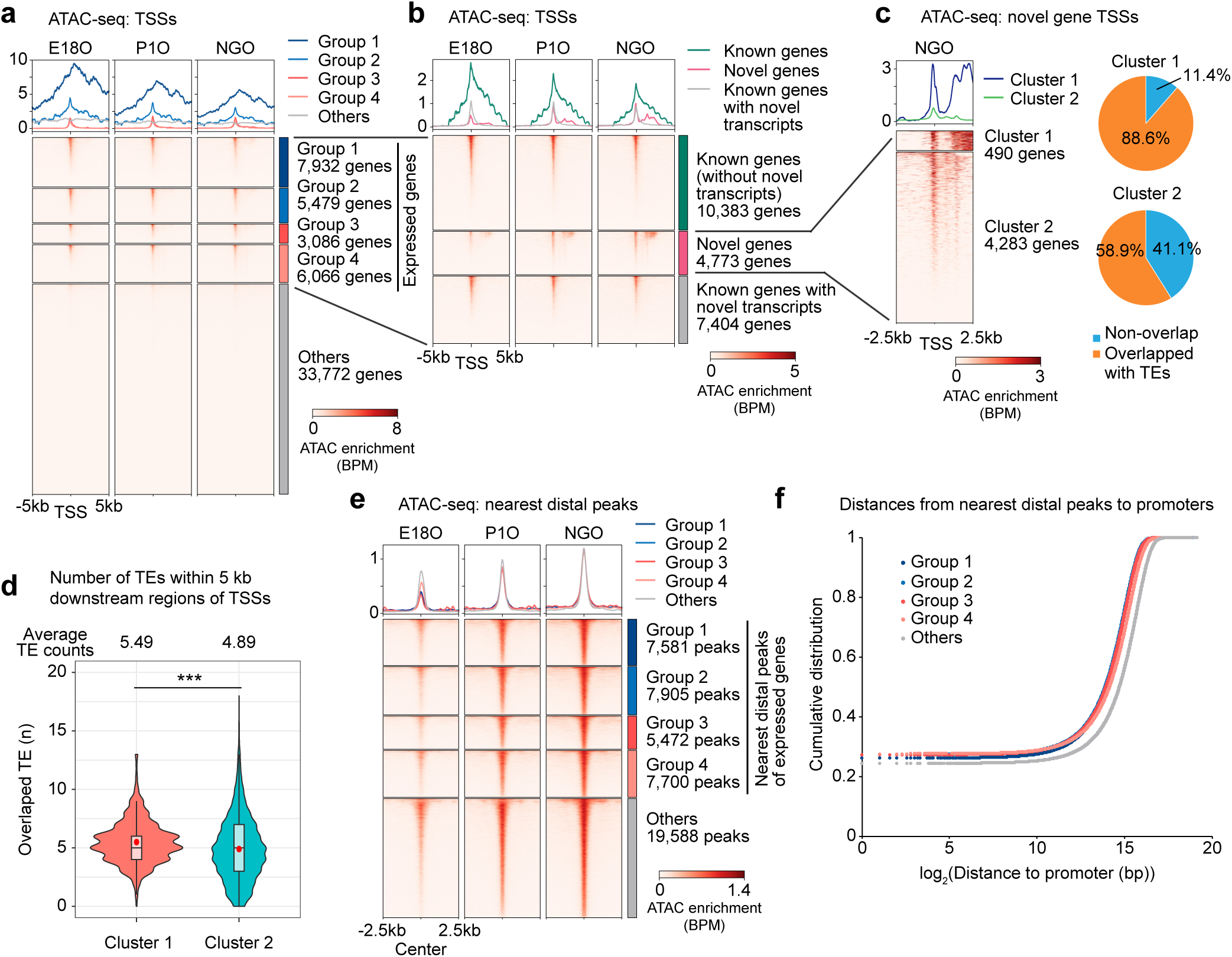
Accessible chromatin dynamics at TSSs and distal regions. **a**, Average tag density plots and heatmaps showing ATAC-seq enrichment dynamics at TSSs for each gene cluster defined in Figure 1e. **b**, Average tag density plots and heatmaps showing ATAC-seq enrichment dynamics at TSSs across all assembled gene annotations. **c**, Average tag density plots and heatmaps showing ATAC-seq enrichment profiles (left), and pie charts illustrating the proportion of genes whose TSSs overlap with TEs (right) among novel genes in NGOs. **d**, Violin plots with overlaid box plots showing the number of TEs located within 5 kb downstream of TSSs in each cluster shown in Figure 3c. The central line indicates the median; box limits represent the 25th and 75th percentiles; whiskers extend to the most extreme data points within 1.5× the interquartile range (IQR). Red dots denote the mean values. \*\*\**P*□<□0.001; Wilcoxon rank-sum test. **e**, Average tag density plots and heatmaps showing ATAC-seq enrichment dynamics at distal peaks (>1 kb from TSSs) for each cluster defined in Figure 1e. **f**, Cumulative distribution plot showing the distances between the promoters of genes in each cluster (Figure 1e) and their nearest distal ATAC-seq peaks.

We next examined the relationship between the gain of distal accessible regions during POT and gene expression. From E18O to NGO, a substantial increase in ATAC signal intensity was observed at the nearest distal peaks across all groups of expressed genes (Fig. 3e), all of which were eventually robustly expressed in GOs (Fig. 1f). Notably, distal accessible regions associated with actively expressed genes (Groups 1–4) were located closer to TSSs than those associated with low or non-expressed genes (Others) (Fig. 3f). Because enhancer activity depends on genomic distance^42^, this observation suggests that the gain of distal accessible regions during POT is linked to the upregulation of nearby genes in GOs, rather than in NGO.

### ERVs serve as proximal and distal regulatory elements that drive the oocyte transcriptome

Having identified associations between TEs and gene expression, we next examined the genome-wide regulation of TEs during perinatal oogenesis. Across the mouse genome, distal accessible regions detected one or more of four representative stages of perinatal oogenesis were relatively enriched for TEs, whereas proximal accessible regions showed little selectivity for TEs compared to the expected overlaps based on their genomic proportion (Fig. 4a). Notably, both proximal and distal accessible chromatin regions overlapping with TEs were more enriched for the ERV family compared to other TE classes, whereas LINEs were underrepresented (Fig. 4b). Among all ERV types (489 in total) in three ERV familes, we identified 62 ERV types enriched in accessible chromatin during perinatal oogenesis, comprising 46 ERVK types, 4 ERVL types, and 12 ERVL-MaLR types (Fig. 4c). Each distal ERV locus belonging to these ERV types became progressively accessible during POT (Fig. 4d). In addition to these distal loci, proximal ERV loci belonging to the ERVL-MaLR family (e.g., MTA and MTB types) also gained accessibility during POT (Fig. 4e), and this gain of accessibility in MTA and MTB types preceded their robust expression in GOs (Fig. 4f, g). In contrast, the mouse-specific ERVK family RLTR types (specifically RLTR10, RLTR10-int, and RLTR10A) were already accessible from E18O onward and were highly upregulated in GOs (Fig. 4f, g). Together, these analyses define a dynamic landscape of TE accessibility during perinatal oogenesis, highlighting ERV subfamilies that undergo stage-specific chromatin activation.

**Figure 4.**
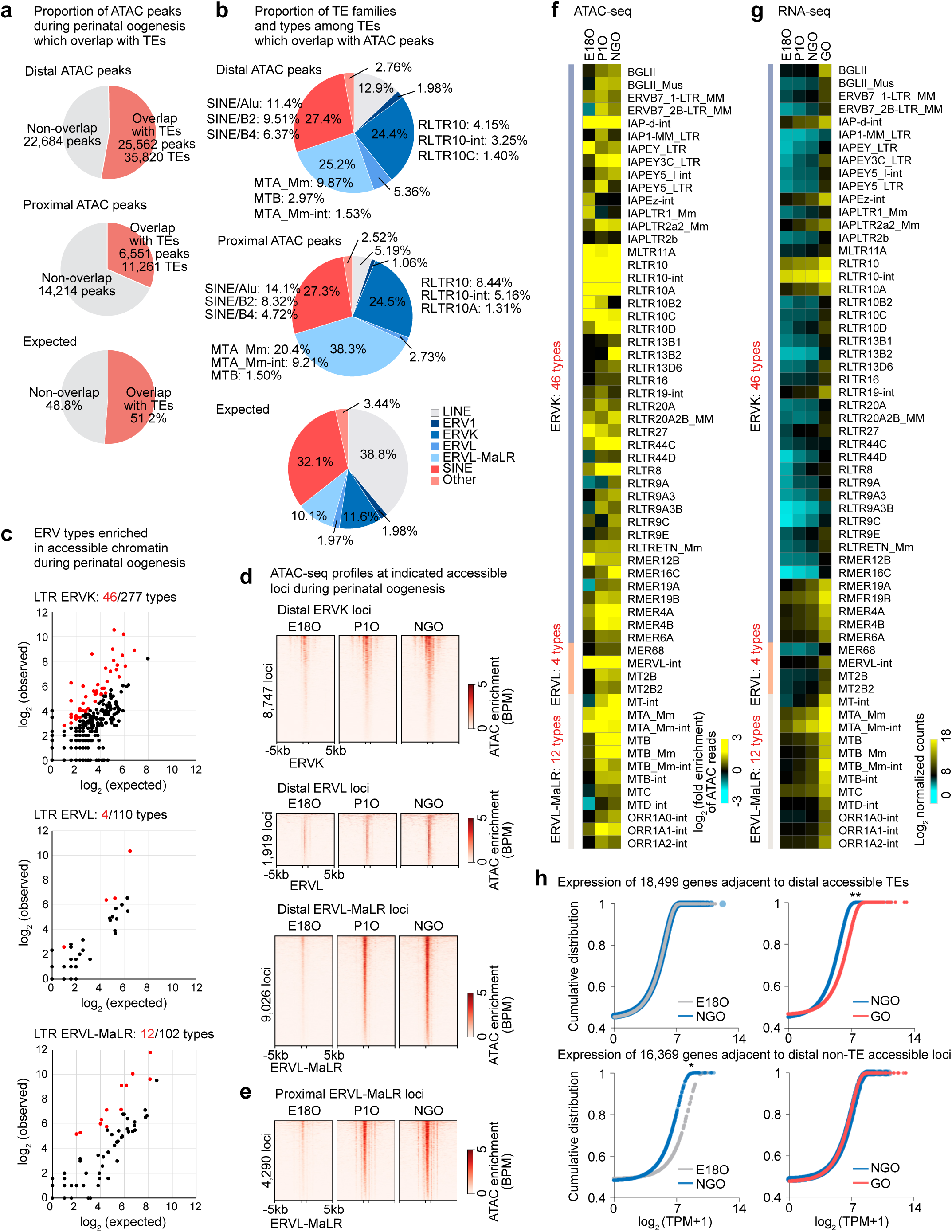
Identification of ERVs in accessible chromatin regions. **a,** Pie charts showing the proportion of ATAC-seq peaks during perinatal oogenesis that overlap with TEs in distal (top) and proximal (middle) regions, and the expected proportion of overlaps based on the genomic TE composition (bottom). **b,** Pie charts showing the distribution of TE families and types among TEs overlapping ATAC-seq peaks in distal (top) and proximal (middle) regions, and the expected distribution based on the genomic prevalence of each TE family (bottom). **c,** Scatter plots showing observed ATAC-seq read counts for each ERV type within accessible chromatin regions (y-axis) versus their expected genomic prevalence (x-axis) for the ERVK, ERVL, and ERVL– MaLR families in NGOs. Each dot represents a single ERV type within a subfamily; red dots indicate ERV types significantly enriched in ATAC-seq reads (≥2-fold observed/expected enrichment; *P*□<□0.05, binomial test). **d,** Heatmaps showing ATAC-seq enrichment at accessible distal ERV regions during POT. **e,** Heatmaps showing ATAC-seq enrichment at accessible proximal ERVL–MaLR regions during POT. **f,** Heatmaps showing ATAC-seq enrichment at ERV types enriched in accessible chromatin during perinatal oogenesis. **g,** Heatmaps showing ERCC-normalized expression levels of ERV types enriched in accessible chromatin during perinatal oogenesis. **h,** Cumulative distribution plots comparing the expression of genes during POT: 18,499 genes adjacent to distal accessible TEs (top) and 16,369 genes adjacent to distal non-TE accessible regions (bottom) at E18O (gray), NGO (blue), and GO (red). \*\**P* < 0.01; \**P* < 0.05; two-tailed unpaired *t*-tests.

Importantly, genes adjacent to distal accessible TE loci during perinatal oogenesis were highly expressed in GOs compared to genes adjacent to distal non-TE accessible loci (Fig. 4h, right). This observation suggests that distal accessible TE loci established in NGOs act as regulatory elements that drive gene expression in GOs. Interestingly, genes adjacent to distal non-TE accessible loci were downregulated during POT (Fig. 4h, bottom left), raising the possibility that non-TE-mediated gene expression is repressed during POT and shifts toward TE-dependent regulation during PPT. Together, these findings suggest that newly evolved ERVs gain accessibility during POT at both distal and proximal genomic regions, where they are co-opted as regulatory elements that expand the repertoire of gene regulation during perinatal oogenesis.

### PRC1 maintains poised enhancer and promoters in NGOs

Our results indicate that the gain of accessibility at gene regulatory elements, including ERV loci, in NGOs precedes the robust gene expression observed in GOs. Accordingly, we reanalyzed the growing oocytes ATAC-seq data^43^ and observed that ATAC-seq signal intensities in NGOs and GOs were positively correlated (Fig. 5a), and that the majority of accessible regions gained during NGO were maintained in GOs (Fig. 5b), suggesting that these regulatory elements persist in an accessible state. We therefore defined accessible chromatin regions shared between NGO and GO as primed accessible chromatin, poised for subsequent gene activation, and found that these regions were enriched not only at promoters but also within distal regions outside promoters (Fig. 5c).

**Figure 5.**
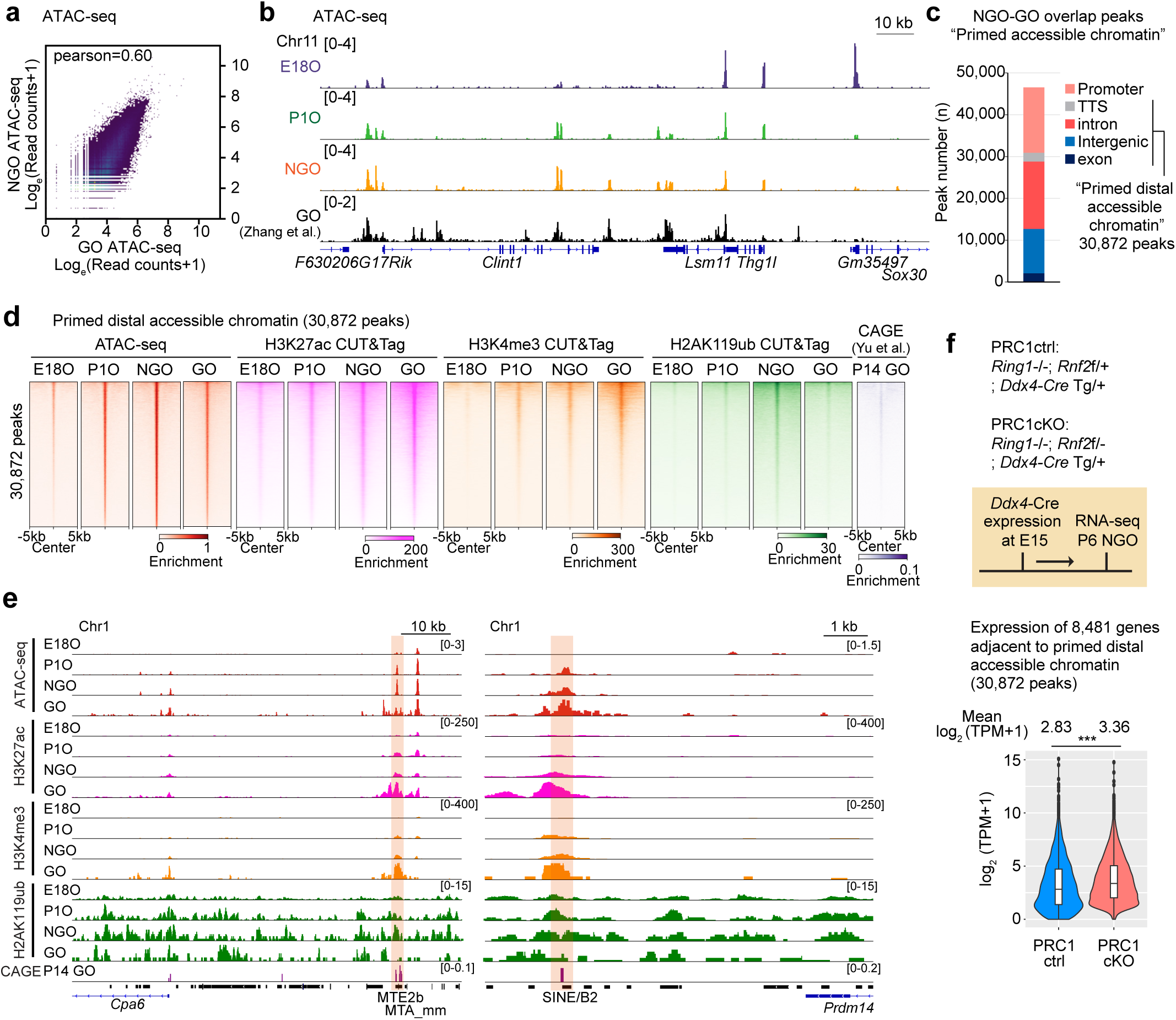
PRC1-poised enhancers in NGO. **a**, Scatter plots showing ATAC-seq enrichment comparing NGO and GO. The Pearson correlation coefficient is indicated. **b**, Track views of ATAC-seq enrichment during perinatal oogenesis. Data ranges are shown in brackets. **c**, Numbers and genomic distribution of ATAC-seq peaks shared between NGO and GO, defined as primed accessible chromatin. **d**, Average tag density plots and heatmaps showing enrichment of ATAC-seq, H3K27ac, H3K4me3, and H2AK119ub during perinatal oogenesis, and CAGE-seq in P14 GO at primed distal accessible chromatin. **e**, Representative track views of ATAC-seq, H3K27ac, H3K4me3, and H2AK119ub enrichment during perinatal oogenesis. Poised enhancer regions within transposable elements are highlighted. Data ranges are shown in brackets. **f**, Schematic of mouse models and RNA-seq experiments (top). Violin plots with a box plot showing RNA expression levels of genes adjacent to primed distal accessible chromatin in PRC1 control (ctrl) and conditional knockout (cKO) NGO (bottom). The central lines represent medians; the boxes represent the interquartile range (IQR; 25th–75th percentile); whiskers extend up to 1.5× IQR from the hinges. \*\*\**P* < 0.001, Wilcoxon rank-sum test.

These results raise the question of how primed accessible chromatin regions are established in NGOs. In oocytes, putative active enhancers are marked not only by H3K27ac enrichment but also by H3K4me3^26^. We therefore investigated whether these histone modifications are already established at primed distal accessible chromatin in NGOs prior to gene activation. By reanalyzing our recent CUT&Tag data^25^, we found that these regions gradually acquired the enhancer marks H3K27ac and H3K4me3 during POT and became strongly enriched in GOs (Fig. 5d). Furthermore, CAGE-seq analysis^44^ revealed the transcription of RNAs—presumably enhancer RNAs—from these regions in GOs (Fig. 5d), suggesting that primed accessible chromatin regions are poised to function as active enhancers in GOs.

To investigate how primed accessible chromatin regions remain repressed in NGOs until follicle activation, we focused on H2A119ub, a repressive histone modification mediated by PRC1, which plays a crucial role in epigenetic programming during POT^23^. Consistent with the high levels of H2A119ub in NGOs^25^, we observed strong enrichment of H2A119ub at primed distal accessible chromatin regions, which was lost during PPT (Fig. 5d, e). Furthermore, TE-derived primed accessible promoters are also marked by H2A119ub (Extended Data Fig. 5a). To determine whether PRC1 suppresses the activity of these regions in NGOs, we analyzed the expression of genes adjacent to primed distal accessible chromatin regions in germ cell–specific PRC1 knockout (PRC1-cKO) mice, in which PRC1 function was eliminated after embryonic day 15 (E15) using *Ddx4*-Cre to mutate the E3 ubiquitin ligase RNF2 on a background lacking its partially redundant paralog RING1 (RING1A)^23^. Compared to PRC1 control (PRC1-ctrl) mice, which lack RING1 but show normal oogenesis^23^, these genes were derepressed in NGOs (Fig. 5f). Together, these findings suggest that primed accessible chromatin regions are established as potential enhancers in NGOs but repressed by PRC1 until follicle activation during PPT.

### TCF3 and TCF12 are already loaded in NGOs to prepare for transcriptional activation

TCF3 and TCF12 are key transcription factors driving the large-scale gene activation associated with PPT^26^. Consistent with this function, our HOMER motif analysis identified TCF3 and TCF12 binding motifs in both proximal and distal primed regions (Fig. 6a). Further analysis revealed that 45% of primed accessible chromatin regions overlapped with TEs, which were particularly enriched for LTR elements from the ERVK and ERVL-MaLR families (Fig. 6b, c). Indeed, accessible ERVs —including MTA and RLTR10 elements—harbor bHLH transcription factor binding motif, such as those for the TCF3 and TCF12 motifs (consensus sequence: CAGCTG; Extended Data Fig. 5b), suggesting that the acquisition of these TEs may have rewired the gene regulatory network to drive transcriptional activation upon follicle activation.

**Figure 6.**
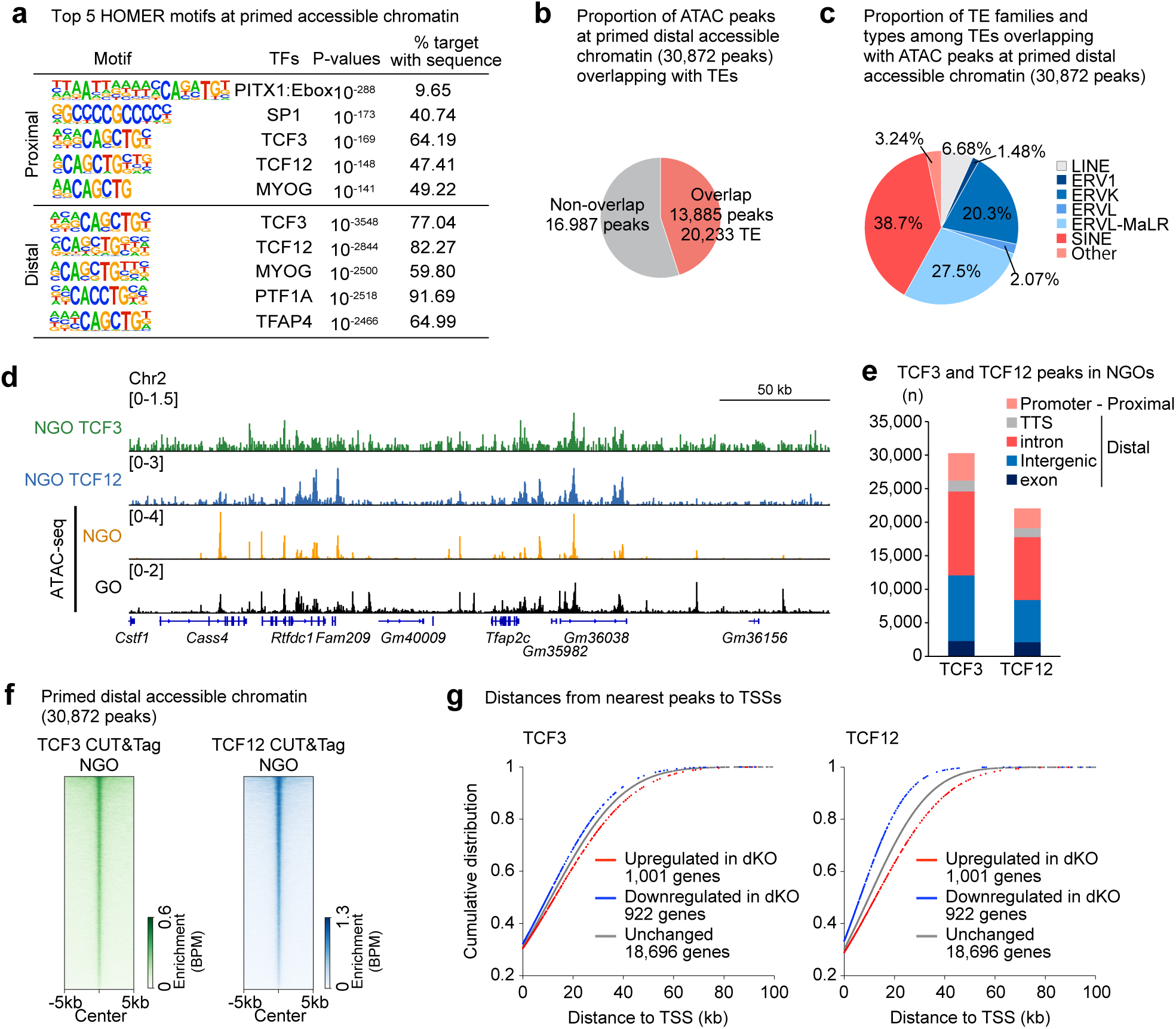
TCF3 and TCF12 are already loaded in NGO. **a**, HOMER known motif analyses of ATAC-seq proximal (top) and distal (bottom) peaks shared between NGO and GO. **b**, Pie charts showing the proportion of ATAC-seq peaks at primed distal accessible chromatin overlapping with TEs. **c**, Pie charts showing the proportion of TE families and types among TEs overlapping with ATAC-seq peaks at primed distal accessible chromatin. **d**, Track views of TCF3 and TCF12 enrichment and ATAC-seq enrichment in NGO and GO. **e**, Numbers and genomic distribution of TCF3 and TCF12 peaks in NGO. **f**, Heatmaps showing TCF3 and TCF12 enrichment in NGO at distal ATAC-seq peaks shared between NGO and GO. **g**, Cumulative distribution plots comparing the distance between TSSs of upregulated, downregulated, and unchanged genes in *Tcf3*/*Tcf12* double-knockout (dKO) GOs and the nearest TCF3 (left) or TCF12 (right) peaks in NGO. Dysregulated genes (1,100 upregulated and 922 downregulated in Tcf3/Tcf12 dKO GOs) were identified by reanalyzing the RNA-seq data of Tcf3/Tcf12 dKO GOs (Ref. ^26^) using the following criteria: fold change ≥ 2 and adjusted *P* < 0.05 (binomial test with Benjamini–Hochberg correction).

To determine whether TCF3 and TCF12 are bound to these regions prior to follicle activation— similar to the pattern observed for H3K27ac—we performed CUT&Tag analysis^45^ for TCF3 and TCF12 in NGOs, identifying binding peaks by merging independent biological replicates (Extended Data Fig. 6a). The resulting CUT&Tag peaks displayed a distribution pattern similar to that of ATAC-seq peaks in both NGO and GO (Fig. 6d). The majority of TCF3 and TCF12 binding sites were enriched in intronic and intergenic regions, rather than promoter regions (Fig. 6e). Notably, binding was also observed at accessible chromatin in promoter-proximal regions in addition to distal sites (Extended Data Fig. 6b). Motif analysis further confirmed that these CUT&Tag peaks contain consensus binding motifs for TCF3, TCF12, and other bHLH transcription factors (Extended Data Fig. 6c).

Importantly, TCF3 and TCF12 were enriched at primed distal accessible chromatin regions shared between NGOs and GOs (Fig. 6f). These binding peaks were located near genes that were downregulated in GOs of oocyte-specific *Tcf3/Tcf12* double-knockout (dKO) mice^26^, in which *Tcf3* and *Tcf12* were deleted in NGOs using *Gdf9*-Cre (Fig. 6g). These results indicate that TCF3 and TCF12 upregulate expression of neighboring genes. Together, these findings suggest that TCF3 and TCF12 occupy gene regulatory elements in advance to prime them for subsequent activation.

### Non-phosphorylated Pol II primes regulatory elements in advance

To enable rapid transcriptional responses during development, RNA polymerase II (Pol II) is often recruited to promoter regions and held in a paused state near TSSs through phosphorylation at serine 5 (Ser5) of its C-terminal domain (CTD), thereby priming genes for activation^46^. To examine whether Pol II is recruited to primed accessible chromatin regions during perinatal oogenesis, we performed CUT&Tag analysis in E18O, NGO, and GO using antibodies against total Pol II (which also detect the non-phosphorylated form), Ser5-phosphorylated Pol II (Ser5P), and Ser2-phosphorylated Pol II (Ser2P), the latter marking elongating Pol II.

Interestingly, total Pol II exhibited two major transitions—one between E18O and NGO (POT), and another from NGO to GO (PPT). In contrast, changes in Ser5P and Ser2P Pol II were detected only between NGOs and GOs (Fig. 7a, Extended Data Fig. 7a). These findings suggest that, rather than being paused via Ser5 phosphorylation, Pol II is recruited to regulatory regions in a non-phosphorylated state in NGOs, potentially enabling long-term chromatin priming prior to transcriptional activation in GOs.

**Figure 7.**
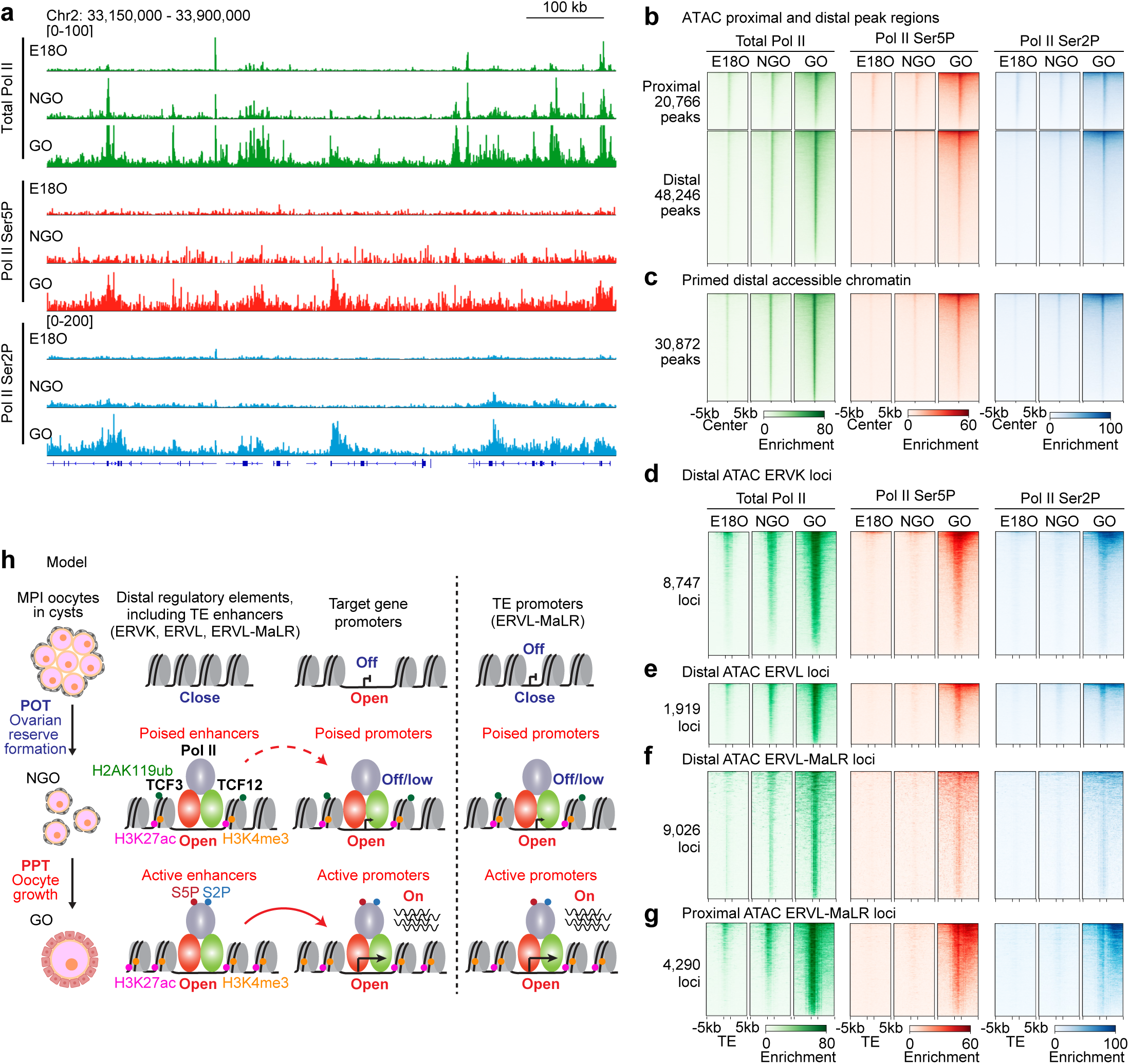
Dynamics of Pol II accumulation during perinatal oogenesis. **a**, Track views of total Pol II, Pol II Ser5P and Pol II Ser2P enrichment during perinatal oogenesis. **b**, Heatmaps showing total Pol II, Pol II Ser5P, and Pol II Ser2P enrichment dynamics at proximal (TSS ±1 kb) and distal (>1 kb from TSSs) peaks during perinatal oogenesis. **c**, Heatmaps showing total Pol II, Pol II Ser5P, and Pol II Ser2P enrichment dynamics at distal ATAC-seq peaks shared between NGO and GO during perinatal oogenesis. **d–f**, Heatmaps showing total Pol II, Pol II Ser5P, and Pol II Ser2P enrichment dynamics at accessible distal ERVK (d), ERVL (e), and ERVL-MaLR (f) regions during perinatal oogenesis. **g**, Heatmaps showing total Pol II, Pol II Ser5P, and Pol II Ser2P enrichment dynamics at accessible proximal ERVL-MaLR regions during perinatal oogenesis. **h**, Schematic model of poised enhancers and promoters primed for oocyte growth.

Total Pol II was recruited to both proximal and distal accessible chromatin regions in NGOs and further enriched in GOs, while Ser5P and Ser2P Pol II were strongly enriched only in GOs (Fig. 7b). A similar trend was observed at primed distal accessible chromatin regions shared between NGOs and GOs (Fig. 7c), indicating that non-phosphorylated Pol II is associated with chromatin priming in NGOs. We also examined Pol II behavior at distal and proximal ERV loci that acquire de novo accessibility. These loci similarly recruited total Pol II in NGOs, with further enrichment in GOs, whereas Ser5P and Ser2P Pol II were enriched only in GOs (Fig. 7d–g). Together, these results indicate that Pol II is already loaded onto primed accessible chromatin regions in NGOs, not in the typical paused (Ser5P) or elongating (Ser2P) forms, but instead in a non-phosphorylated, poised state. In addition, the presence of Ser5P and Ser2P on the novel genes identified in this study corroborates their transcription in GOs (Extended Data Fig. 7b, c).

In summary, we demonstrate that the large-scale transcriptional transition that occurs during perinatal oogenesis is driven by the genome-wide establishment of accessible chromatin, particularly at newly evolved TEs, which function as poised enhancers and promoters in preparation for follicular activation. These primed regions are already bound by transcription factors such as TCF3 and TCF12, along with non-phosphorylated Pol II, thereby establishing a poised transcriptional landscape in NGOs that is stably maintained until gene activation in GOs.

## Discussion

In this study, we redefine the oocyte transcriptome during perinatal oogenesis and uncover a vast repertoire of previously unannotated transcripts whose expression is driven by newly evolved ERVs. We demonstrate that accessible chromatin regions—particularly those enriched in TEs—are established during ovarian reserve formation and primed for genome-wide transcriptional activation at PPT (Fig. 7h). Notably, putative enhancer regions in GOs are already established and primed in NGOs, indicating that chromatin states are preconfigured and stably maintained within the ovarian reserve. Thus, our findings identify epigenetic priming in NGOs as a key prerequisite for the transition from quiescence to growth, revealing a previously unrecognized mode of chromatin-based preparation for follicle activation.

Building on previous studies showing that ERVL-MaLR elements regulate a subset of the oocyte transcriptome^19–22^, we identified novel transcripts with TE-derived promoters not only in GOs but also in MPI oocytes. Notably, promoter usage shifted from ERVK (RLTR10), which were accessible in E18O to NGO, to ERVL-MaLR (MTA), which gained accessibility during POT. This suggests a shift in the cellular transcriptomes driven by TEs during perinatal oogenesis (Fig. 1h). We further observed a genome-wide increase in chromatin accessibility, especially at distal regions enriched for ERVK (RLTR10) and ERVL-MaLR (MTA), during ovarian reserve formation, i.e., prior to follicular activation. These findings suggest that both proximal and distal ERVs play instructive roles in shaping the oocyte transcriptome. These results corroborate previous findings that mouse-specific MaLR elements, such as MTA, serve as promoters in mouse oocytes^21,22^. Importantly, both RLTR10 and MTA elements harbor bHLH transcription factor binding motifs, including sites for TCF3 and TCF12. The acquisition of these newly evolved ERVs appears to have redistributed gene regulatory elements across the genome, thereby contributing to the epigenetic priming that enables follicular activation and defines the subsequent oocyte transcriptome.

In the broader context of germline biology, TE-driven follicular activation shares similarities with TE-mediated gene expression during zygotic genome activation (ZGA) following fertilization^47,48^. In the mouse zygote, expression of mouse-specific ERVL elements during ZGA is essential for proper preimplantation development^49^. This parallel suggests a shared evolutionary strategy, whereby the acquisition of newly evolved TEs enhances reproductive fitness by shaping genome-wide transcriptional programs during both follicular activation and ZGA. Moreover, a similar wave of genome-wide transcriptional activation occurs during male meiosis, where RLTR10 elements function as distal enhancers^50^. This finding implies a potential parallel between male and female MPI, further underscoring the notion that TE activity regulates germline-specific gene expression. Importantly, our findings show that TEs encode a chromatin-based memory that contributes to epigenetic priming in oocytes. By establishing accessible chromatin and recruiting the transcriptional machinery prior to gene activation, TEs likely facilitate rapid transcriptional activation upon follicle activation.

Epigenetic priming involves both activating and repressive histone modifications that prepare chromatin for rapid gene activation. Traditionally, bivalent histone marks—co-enrichment of H3K4me3 and H3K27me3—have been implicated in gene regulation during stem cell differentiation and germline development^51–54^. In this study, we identify a distinct poised chromatin state in NGOs, marked by the co-localization of H3K27ac and H2AK119ub at both distal accessible regions and proximal promoters. Notably, PRC2-mediated H3K27me3, which typically antagonizes H3K27ac, appears to play a minor role in NGOs, but H3K27me3 increases later in GOs^25^. These findings suggest that, unlike the canonical H3K4me3/H3K27me3 bivalent domains regulated by PRC2, epigenetic priming in NGOs is predominantly governed by PRC1-mediated repression. This proposal is supported by previous reports showing that oocyte-specific deletion of PRC1 leads to gene de-repression and premature depletion of the ovarian reserve, underscoring the critical role of PRC1 in maintaining transcriptional quiescence and oocyte longevity^23^.

Promoter-proximal pausing of Pol II, marked by phosphorylation of Ser5 on its CTD, is a recognized feature of epigenetic priming that positions the transcriptional machinery for rapid activation during developmental transitions^46,55^. In NGOs, we find that Pol II is already loaded at both proximal and distal regulatory elements in an unphosphorylated state (Fig. 7). Given the long-term quiescence of NGOs, this unphosphorylated Pol II may represent a stable form of transcriptional poising that supports the longevity of the primed state.

Together, our findings uncover a non-canonical chromatin priming mechanism in NGOs, driven by PRC1 activity and unphosphorylated Pol II loading, which prepares the oocyte genome for rapid activation upon growth initiation. This study provides a foundation for future investigations into the regulatory mechanisms governing follicular activation, a largely unexplored area of gene regulation in the germline.

## Acknowledgments

We thank members of the Namekawa lab and Neil Hunter for the discussion; Azim Surani for providing *Stella*-GFP transgenic mice; and So Maezawa for the homemade Tn5 transposase for ATAC-seq. We acknowledge the following funding sources: JSPS Overseas Challenge Program for Young Researchers, JSPS Overseas Research Fellowship, Lalor foundation postdoctoral fellowship, Productive Health Global Consortium (PHGC) postdoctoral fellowship 2223 to Y.M.; Repro Grant to M. H.; JSPS KAKENHI Grant JP23K23898 to N.I.; and NIH Grants R35GM141085 to S.H.N and R21HD110146 to R.M.S. and S.H.N.

## Author contributions

Y.M. and S.H.N. designed the study. Y.M. performed the RNA-seq, ATAC-seq and CUT&Tag experiments, with the contribution from M.H. Y.M. performed the computational analyses, with the contribution from R.D. All authors interpreted the data. Y.M., R.M.S., and S.H.N wrote the manuscript with critical feedback from all authors. S.H.N. supervised the project.

## Methods

### Animals

Mice were maintained on a 12:12 light cycle in a temperature and humidity-controlled vivarium (22±2°C: 40-50% humidity) with free access to food and water in a pathogen-free animal care facility. Mice were used according to the guidelines of the Institutional Animal Care and Use Committee (protocol no. IACUC2018-0040, 21943, and 23545) at Cincinnati Children’s Hospital Medical Center and the University of California, Davis. *Stella*-GFP transgenic mice were obtained from Dr M. Azim Surani,^56^ maintained on a mixed genetic background of FVB and C57BL/6J.

### Oocyte collection

Female pups were collected at different developmental stages, and ovaries were harvested by carefully removing oviducts and ovarian bursa in phosphate-buffered saline (PBS). Ovaries were digested in 200 μl TrypLE™ Express Enzyme (1X) (Gibco, 12604013) supplemented with 0.3% Collagenase Type 1 (Worthington, CLS-1) and 0.01% DNase I (Sigma, D5025) and incubated at 37°C for 30 min with gentle agitation. After incubation, the ovaries were dissociated by gentle pipetting using the FisherbrandTM Premium Plus MultiFlex Gel-Loading Tips until no visible tissue pieces. Two ml Leibovitz’s L-15 Medium (Gibco, 11415064) supplemented with 10% FBS (HyClone, SH30396.03) were then added to the suspension to stop enzyme reaction.

For FACS preparation, the cells were then washed with FACS buffer (PBS containing 2%FBS) twice by centrifugation at 300 × g for 5 min and filtered into a 5 ml FACS tube with a 35 μm nylon mesh cap (Falcon, 352235). *Stella*-GFP^+^ oocytes were collected after removing small and large debris in FSC-A versus SSC-A gating and doublets in FSC-W versus FSC-H gating.

For manual picking, the cell suspension was filtered through a 100 μm cell strainer and seeded onto a culture dish. The cells were allowed to settle down for 15-30 min at 37°C before being transferred under the stereomicroscope (Nikon, SMZ1270). Based on morphology and diameter, growing oocytes were specifically picked up, washed in M2 medium (Sigma, M7167), and transferred into the downstream buffer by mouth pipette with a glass capillary.

### Strand-specific total RNA-seq library generation with ERCC RNA spike-in and sequencing

RNA-seq libraries of oocytes were prepared as follows. The non-growing oocytes isolated from ovaries at E18.5, P1, and P5-6 using FACS, and the growing oocytes isolated from P6-7 ovaries under the stereomicroscope. Two independent biological replicates were used for RNA-seq library generation. Total RNA was extracted using the RNeasy Plus Micro Kit (QIAGEN, Cat # 74034) according to the manufacturer’s instructions and the External RNA Controls Consortium (ERCC) RNA Spike-In Mix (Invitrogen, 4456740) was added to total RNA. This mix was first diluted 100-fold with nuclease-free water, and then 1 µL of this dilution was added to extracted total RNA from 10,000 cells. Ribosomal RNA was depleted from total RNA using NEBNext® rRNA Depletion Kit v2 (Human/Mouse/Rat) (NEB, E7400L). Library preparation was performed with NEBNext® Ultra™ II Directional RNA Library Prep Kit for Illumina® (NEB, E7760S) according to the manufacturer’s instruction. Prepared RNA-seq libraries were sequenced on the NovaSeq X Plus system (Illumina) with paired-ended 150-bp reads, and each stage was profiled in three biological replicates. In total, we generated 1.35 billion reads, of which 1.12 billion were uniquely mapped concordant paired-end reads.

### ATAC-seq library generation and sequencing

ATAC-seq libraries of oocytes were prepared as described^38^. Briefly, 10,000 FACS-sorted oocytes were isolated from ovaries at E18.5, P1, and P5-6 and pooled as one replicate each different developmental stages, and two independent biological replicates were used for ATAC-seq library generation. oocytes were lysed in 50 μl of lysis buffer [10 mM Tris–HCl (pH 7.4), 10 mM NaCl, 3 mM MgCl2 and 0.1% NP-40, 0.1% Tween-20, and 0.01% Digitonin] on ice for 10 min. Immediately after lysis, the samples were spun at 500 × g for 10 min at 4°C and the supernatant removed. The sedimented nuclei were then incubated in 10 μl of transposition mix [0.5 μl homemade Tn5 transposase (∼1 μg/μl), 5 μl 2× tagmentation DNA buffer [10 mM Tris–HCl (pH 7.6), 10 mM MgCl2 and 20% dimethyl formamide], 3.3 μl PBS, 0.1 μl 1% digitonin, 0.1 μl 10% Tween-20 and 1 μl water] at 37°C for 30 min in a thermomixer with shaking at 500 rpm. After tagmentation, the transposed DNA was purified with a MinElute kit (Qiagen). Polymerase chain reaction (PCR) was performed to amplify the library using the following conditions: 72°C for 3 min; 98°C for 30 s; thermocycling at 98°C for 10 s, 60°C for 30 s and 72°C for 1 min. Quantitative PCR was used to estimate the number of additional cycles needed to generate products at 25% saturation. Seven to eight additional PCR cycles were added to the initial set of five cycles. Amplified DNA was purified by SPRIselect bead (Beckman Coulter). ATAC-seq libraries were sequenced on the HiSeq X system (Illumina) with 150-bp paired-end reads.

### Quantitative CUT&Tag library generation and sequencing

CUT&Tag libraries of oocytes were prepared as previously described^45,57^ with some modifications (a step-by-step protocol https://www.protocols.io/view/bench-top-cut-amp-tag-kqdg34qdpl25/v3) using CUTANA™ pAG-Tn5 (Epicypher, 15-1017). To perform quantitative spike-in CUT&Tag, Drosophila S2 cells were added to mouse oocytes at a fixed ratio (e.g., 5,000 S2 cells to 10,000 mouse oocytes) at the beginning of each reaction. The antibodies used were mouse anti-E2A (G-2) (TCF3, 1/50; Santa Cruz Biotechnology; sc-133075), mouse anti-HEB (A-6) (TCF12, 1/50; Santa Cruz Biotechnology; sc-365980), rabbit anti-H3K27me3 (1/50; Cell Signaling Technology; 9733), mouse anti-Rpb1 CTD (4H8) (1/50; Cell Signaling Technology; 2629), rabbit anti-Phospho-Rpb1 CTD (Ser5) (D9N5I) (1/50; Cell Signaling Technology; 13523) and r rabbit anti-Phospho-Rpb1 CTD (Ser2) (E1Z3G) (1/50; Cell Signaling Technology; 13499). CUT&Tag libraries were sequenced on the NovaSeq X Plus system (Illumina) with 150-bp paired-end reads.

### RNA-seq data processing

Raw paired-end RNA-seq reads after trimming by Trim-galore (https://github.com/FelixKrueger/TrimGalore) (version 0.6.7) were aligned to the mouse (GRCm38/mm10) genome using STAR (version STAR_2.5.4b)^58^ with following options: --outSAMtype BAM SortedByCoordinate; --twopassMode Basic; --outFilterType BySJout; --outFilterMultimapNmax 1; --winAnchorMultimapNmax 50; --alignSJoverhangMin 8; --alignSJDBoverhangMin 1; -- outFilterMismatchNmax 999; --outFilterMismatchNoverReadLmax 0.04; --alignIntronMin 20; -- alignIntronMax 1000000; --alignMatesGapMax 1000000 for unique alignments. To quantify aligned reads in RNA-seq, aligned read counts for each gene were generated using featureCounts (v2.0.1), which is part of the Subread package^59^ based on annotated genes (GENCODE vM25: GRCm38.p6). The transcripts per million (TPM) values of each gene were for comparative expression analyses and computing the Pearson correlation coefficient between biological replicates using corrplot^60^.

To generate novel oocyte transcript annotation, de novo transcriptome assemblies from strand-specific RNA-seq libraries were generated using StringTie2 (v2.1.2), a reference-guided assembler known for its high accuracy and sensitivity in transcript detection^61,62^, with default parameters. The assembled files and GENCODE reference annotation (Release M25: GRCm38.p6) were merged to generate a non-redundant set of transcripts using StringTie2 with --merge and -G option. Mono-exonic transcripts transcribed from known TE were removed from merged transcript annotation. Repetitive annotation in mm10 genome (mm10.fa.out, open-4.0.5, GRCm38/mm10) was downloaded from the RepeatMasker website (http://www.repeatmasker.org/species/mm.html). TE annotation in this file does not include overlapping. To prepare the TE annotation set in distal regions, TE copies that overlapped with exonic regions of a oocyte transcript annotation or had low SW scores (≤500 for SINE and DNA transposons and ≤1,000 for other transposons) were removed using the BEDTools (version 2.28.0)^63^ intersect function and custom Python scripts. To detect the novel transcript, The novel oocyte transcript annotatio s were compared with GENCODE reference annotations (Release M25: GRCm38.p6) or previously published dataset from Veselovska et al. using Gffcompare (version 0.12.6) (https://doi.org/10.12688/f1000research.23297.1) with default settings.

To quantify uniquely aligned reads, raw aligned read counts for each gene were generated using featureCounts (v2.0.1), which is part of the Subread package based on the novel oocyte transcript annotation and ERCC annotated sequences. The aligned read counts for each transcript were generated using the Salmon (1.9.0)^64^ with quant command and default parameters. For ERCC spike-in normalization and analysis, the edgeR^65^ and Limma^66^ packages were employed. Normalization factors were calculated from reads mapped to ERCC annotated sequences, followed by the filtering of raw reads such that there was a minimum of two counts in at least one sample. The filtered reads were assessed for library size using normalization factors prior to voom transformation. GO analyses were performed using the functional annotation clustering tool in Enrichr (https://maayanlab.cloud/Enrichr/)^67^. RNA expression heatmaps were plotted using Morpheus (https://software.broadinstitute.org/morpheus, Broad Institute). To visualize read enrichments over representative genomic loci using the Integrative Genomics Viewer (Broad Institute)^68^, bins per million (BPM) normalized counts data were created from sorted BAM files using ‘bamCoverage’ tool in deepTools.

To detect differentially expressed genes (DEGs) between *Tcf3*/*Tcf12* ctrl and *Tcf3*/*Tcf12* dKO, DESeq2 (version 1.42.1)^69^ was used for differential gene expression analyses with cutoffs ≥2-fold change and binominal tests (*Padj* < 0.05; *P*-values were adjusted for multiple testing using the Benjamini– Hochberg method). *Padj* values were used to determine significantly dysregulated genes.

### Coding region prediction and amino acid sequence homology

Coding region prediction was performed using CodAn (version 1.2)^35^, CPAT (v3.0.4)^34^, and CPC2 (http://cpc2.gao-lab.org/)^33^. For CodAn analysis, the novel transcripts were separated based on strand orientation into positive and negative strands. FASTA file was generated for each strand using Gffread (version 0.12.7)^70^ and then utilized as an input for the three prediction tools under default parameters. In CodAn, we ran both VERT_full and VERT_partial as the predictive model separately. Subsequently, the predicted peptide sequences were extracted and used as an query to perform BLASTp for amino acid sequence homology with default parameters.

### ATAC-seq data processing

Raw paired-end ATAC-seq reads after trimming by Trim-galore (https://github.com/FelixKrueger/TrimGalore) (version 0.6.7) were aligned to the mouse (GRCm38/mm10) genomes using bowtie2 (version 2.3.3.1)^71^ with options: --end-to-end --very-sensitive - -no-mixed --no-discordant -I 10 -X 2000. The aligned reads were filtered to remove alignments mapped to multiple locations by calling grep with the -v option before being subjected to downstream analyses. PCR duplicates were removed using the ‘MarkDuplicates’ command in Picard tools (version 2.23.8) (https://broadinstitute.github.io/picard/, Broad Institute). The aligned reads were shifted +4□bp and - 5□bp for positive and negative strand respectively using ‘alignmentSieve’ tool in deepTools (version 3.3.0)^72^. To compare replicates, Pearson correlation coefficients were calculated and plotted by ‘multiBamSummary bins’ and ‘plot correlation’ functions of deepTools. Biological replicates were pooled for visualization and other analyses after validation of reproducibility. Peak calling was performed using MACS3 (version 3.0.0a7)^73^ with default arguments. We computed the number of overlapping peaks between peak files using BEDtools (version 2.28.0) function intersect. To detect genes adjacent to ATAC-seq peaks, we used the HOMER (version 4.9.1)^74^ function annotatePeaks.pl. Tag density plots and heatmaps for reads enrichments were drawn using ‘computeMatrix’ and ‘plotHeatmap’ tool in deepTools. To visualize ATAC-seq data using the Integrative Genomics Viewer (Broad Institute), BPM normalized counts data were created from sorted BAM files using ‘bamCoverage’ tool in deepTools. To perform functional annotation enrichment of ATAC-seq peaks, we used GREAT tools^75^.

### CUT&Tag data processing

Raw paired-end CUT&Tag reads after trimming by Trim-galore were aligned to the mouse (GRCm38/mm10) genomes using bowtie2 (version 2.3.3.1) with options: --end-to-end --very-sensitive -- no-mixed --no-discordant -I 10 -X 700. The aligned reads were filtered to remove alignments mapped to multiple locations by calling grep with the -v option before being subjected to downstream analyses. D. melanogaster DNA delivered by Drosophila S2 cells was used as spike-in DNA as described^76^. For mapping D. melanogaster spike-in fragments, we also aligned to either the D. melanogaster (dm6) genome using Bowtie2 and used the ‘--no-overlap --no-dovetail’ options to avoid cross-mapping using Bowtie2. PCR duplicates were removed using the ‘MarkDuplicates’ command in Picard tools. Spike-in normalization was implemented using the exogenous scaling factor computed from the dm6 aligned files (scale factors = 10000/spike-in reads). Spike-in normalized genome coverage tracks with 1 bp resolution in BigWig format were generated using ‘bamCoverage’ tool in deepTools with the parameters ‘--binSize 1 --extendReads --samFlagInclude 64 --normalizeUsing BPM --scaleFactor $scale_factor’. Track views were visualized and exported from Integrative Genomics Viewer. Biological replicates were pooled for visualization and other analyses after validation of reproducibility. Peak calling for TCF3 and TCF12 was performed using the Sparse Enrichment Analysis for CUT&RUN (SEACR) (https://seacr.fredhutch.org/)^77^ by selecting the top 1% enriched regions in stringent mode with parameters’-c 0.01 -n norm -m stringent’. To detect genes adjacent to TCF3 and TCF12 peaks, we used the HOMER (version 4.9.1) function annotatePeaks.pl. To identify the enrichment of known motifs, we used the HOMER function findMotifsGenome.pl with default parameters. Tag density plots and heatmaps for reads enrichments were drawn using ‘computeMatrix’ and ‘plotHeatmap’ tool in deepTools.

### Statistics

Statistical methods and *P* values for each plot are listed in the figure legends and/or in the Methods. In brief, all grouped data are represented as mean ± standard deviation (SD). All box-and-whisker plots are represented as center lines (median), box limits [interquartile range (IQR); 25th and 75th percentiles] and whiskers (maximum value not exceeding 1.5× the IQR from the hinge) unless stated otherwise. Statistical significance for pairwise comparisons was determined using two-sided Mann– Whitney U-tests and two-tailed unpaired t-tests. Next-generation sequencing data (RNA-seq, ATAC-seq and CUT&Tag) were based on two or three independent replicates. No statistical methods were used to predetermine sample size in these experiments. Experiments were not randomized, and investigators were not blinded to allocation during experiments and outcome assessments.

