## Extended Data Figures 1-7 for "Newly Evolved Endogenous Retroviruses Prime the Ovarian Reserve for Activation"

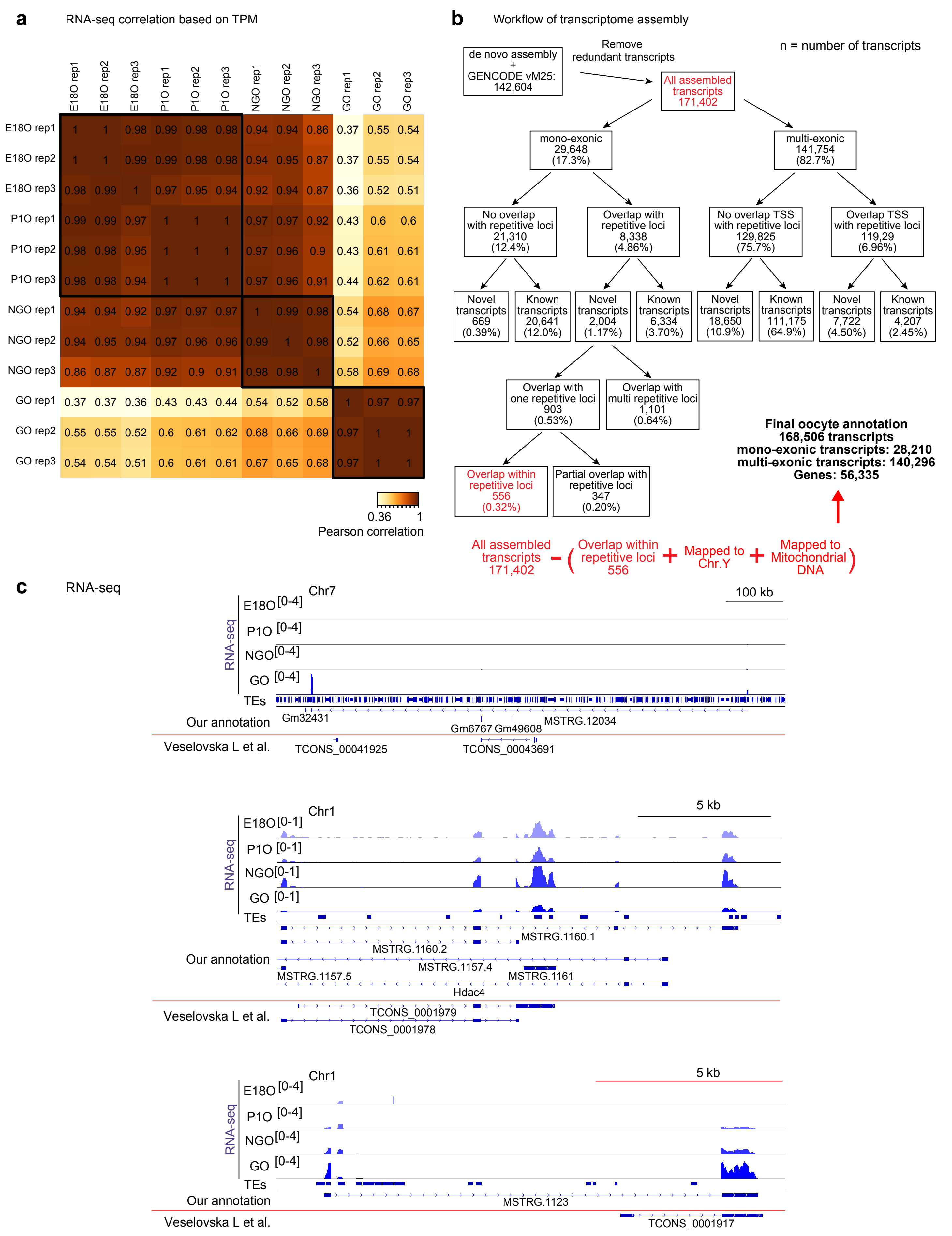
a
RNA-seq correlation based on TPM
E18O rep1
E18O rep2
E18O rep3
P1O rep1
P1O rep2
P1O rep3
NGO rep1
NGO rep2
NGO rep3
GO rep1
GO rep2
GO rep3
E18O rep1 1 1 0.98 0.99 0.98 0.98 0.94 0.94 0.86 0.37 0.55 0.54
E18O rep2 1 1 0.99 0.99 0.98 0.98 0.94 0.95 0.87 0.37 0.55 0.54
E18O rep3 0.98 0.99 1 0.97 0.95 0.94 0.92 0.94 0.87 0.36 0.52 0.51
P1O rep1 0.99 0.99 0.97 1 1 1 0.97 0.97 0.92 0.43 0.6 0.6
P1O rep2 0.98 0.98 0.95 1 1 1 0.97 0.96 0.9 0.43 0.61 0.61
P1O rep3 0.98 0.98 0.94 1 1 1 0.97 0.96 0.91 0.44 0.62 0.61
NGO rep1 0.94 0.94 0.92 0.97 0.97 0.97 1 0.99 0.98 0.54 0.68 0.67
NGO rep2 0.94 0.95 0.94 0.97 0.96 0.96 0.99 1 0.98 0.52 0.66 0.65
NGO rep3 0.86 0.87 0.87 0.92 0.9 0.91 0.98 0.98 1 0.58 0.69 0.68
GO rep1 0.37 0.37 0.36 0.43 0.43 0.44 0.54 0.52 0.58 1 0.97 0.97
GO rep2 0.55 0.55 0.52 0.6 0.61 0.62 0.68 0.66 0.69 0.97 1 1
GO rep3 0.54 0.54 0.51 0.6 0.61 0.61 0.67 0.65 0.68 0.97 1 1
0.36 1
Pearson correlation
b
Workflow of transcriptome assembly
de novo assembly + GENCODE vM25: 142,604
Remove redundant transcripts
n = number of transcripts
All assembled transcripts 171,402
mono-exonic 29,648 (17.3%)
multi-exonic 141,754 (82.7%)
No overlap with repetitive loci 21,310 (12.4%)
Overlap with repetitive loci 8,338 (4.86%)
No overlap TSS with repetitive loci 129,825 (75.7%)
Overlap TSS with repetitive loci 119,29 (6.96%)
Novel transcripts 669 (0.39%)
Known transcripts 20,641 (12.0%)
Novel transcripts 2,004 (1.17%)
Known transcripts 6,334 (3.70%)
Novel transcripts 18,650 (10.9%)
Known transcripts 111,175 (64.9%)
Novel transcripts 7,722 (4.50%)
Known transcripts 4,207 (2.45%)
Overlap with one repetitive loci 903 (0.53%)
Overlap with multi repetitive loci 1,101 (0.64%)
Overlap within repetitive loci 556 (0.32%)
Partial overlap with repetitive loci 347 (0.20%)
Final oocyte annotation
168,506 transcripts
mono-exonic transcripts: 28,210
multi-exonic transcripts: 140,296
Genes: 56,335
All assembled transcripts 171,402 − ( Overlap within repetitive loci 556 + Mapped to Chr.Y + Mapped to Mitochondrial DNA )
c
RNA-seq
Chr7
100 kb
E18O [0-4]
P1O [0-4]
NGO [0-4]
GO [0-4]
TEs
Our annotation
Gm32431
Gm6767 Gm49608
MSTRG.12034
Veselovska L et al.
TCONS_00041925
TCONS_00043691
Chr1
5 kb
E18O [0-1]
P1O [0-1]
NGO [0-1]
GO [0-1]
TEs
Our annotation
MSTRG.1160.1
MSTRG.1160.2
MSTRG.1157.4
MSTRG.1157.5
MSTRG.1161
Hdac4
Veselovska L et al.
TCONS_0001979
TCONS_0001978
Chr1
5 kb
E18O [0-4]
P1O [0-4]
NGO [0-4]
GO [0-4]
TEs
Our annotation
MSTRG.1123
Veselovska L et al.
TCONS_0001917

**Extended Data Figure 1. De novo transcriptome assembly during perinatal oogenesis.**
**a**, Heatmaps showing Pearson correlations of TPM values among biological replicates in strand-specific total RNA-seq datasets. Pearson correlation coefficients are indicated in the panels.
**b**, Flow chart showing the number of transcripts in each category: monoexonic versus multiexonic, known versus novel, and overlapping with repeats versus non-overlapping. Redundant transcripts were removed by merging de novo assemblies with the GENCODE reference annotation. Monoexonic transcripts fully overlapping with RepeatMasker-annotated TEs were excluded to eliminate non-uniquely mappable TE-derived transcripts.
**c**, Representative track views showing RNA-seq signals, the oocyte transcript annotation generated in this study, and the previously published oocyte annotation from Veselovska et al.[20] Data ranges are shown in brackets.

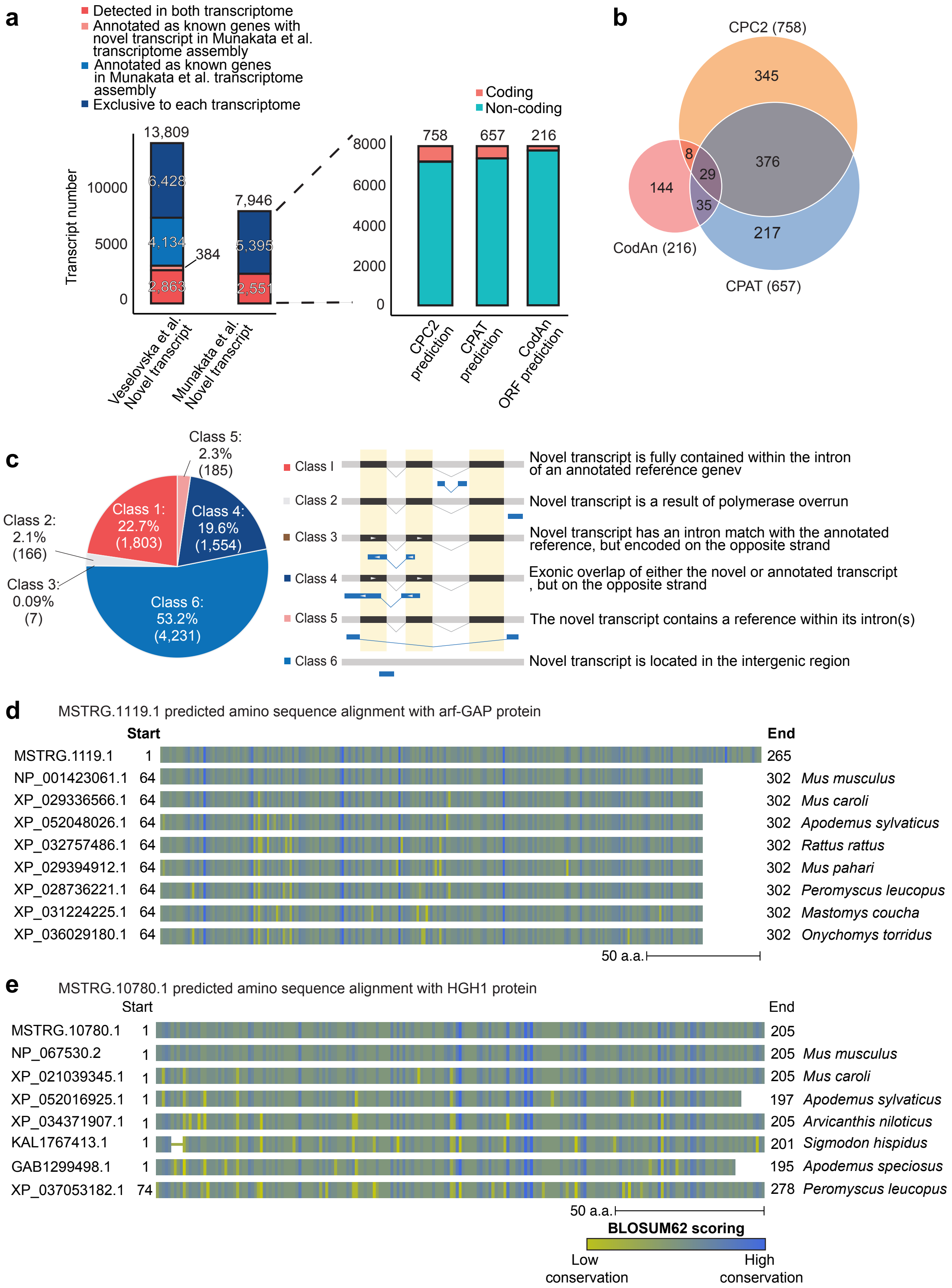
a
Detected in both transcriptome
Annotated as known genes with novel transcript in Munakata et al. transcriptome assembly
Annotated as known genes in Munakata et al. transcriptome assembly
Exclusive to each transcriptome
Transcript number
13,809
6,428
4,134
384
2,863
7,946
5,395
2,551
Veselovska et al. Novel transcript
Munakata et al. Novel transcript
Coding
Non-coding
758
657
216
CPC2 prediction
CPAT prediction
CodAn ORF prediction
b
CPC2 (758)
CPAT (657)
CodAn (216)
345
376
217
144
8
29
35
c
Class 1: 22.7% (1,803)
Class 2: 2.1% (166)
Class 3: 0.09% (7)
Class 4: 19.6% (1,554)
Class 5: 2.3% (185)
Class 6: 53.2% (4,231)
Class I
Novel transcript is fully contained within the intron of an annotated reference genev
Class 2
Novel transcript is a result of polymerase overrun
Class 3
Novel transcript has an intron match with the annotated reference, but encoded on the opposite strand
Class 4
Exonic overlap of either the novel or annotated transcript , but on the opposite strand
Class 5
The novel transcript contains a reference within its intron(s)
Class 6
Novel transcript is located in the intergenic region
d
MSTRG.1119.1 predicted amino sequence alignment with arf-GAP protein
Start
End
MSTRG.1119.1 1 265
NP_001423061.1 64 302 Mus musculus
XP_029336566.1 64 302 Mus caroli
XP_052048026.1 64 302 Apodemus sylvaticus
XP_032757486.1 64 302 Rattus rattus
XP_029394912.1 64 302 Mus pahari
XP_028736221.1 64 302 Peromyscus leucopus
XP_031224225.1 64 302 Mastomys coucha
XP_036029180.1 64 302 Onychomys torridus
50 a.a.
e
MSTRG.10780.1 predicted amino sequence alignment with HGH1 protein
Start
End
MSTRG.10780.1 1 205
NP_067530.2 1 205 Mus musculus
XP_021039345.1 1 205 Mus caroli
XP_052016925.1 1 197 Apodemus sylvaticus
XP_034371907.1 1 205 Arvicanthis niloticus
KAL1767413.1 1 201 Sigmodon hispidus
GAB1299498.1 1 195 Apodemus speciosus
XP_037053182.1 74 278 Peromyscus leucopus
50 a.a.
BLOSUM62 scoring
Low conservation
High conservation

**Extended Data Figure 2. Characterization of novel transcripts.**
**a**, Comparison between novel transcripts identified in Veselovska et al. and in this study (left). Proportions of novel transcripts with coding potential predicted by CPC2, CPAT, and CodAn (right).
**b**, Venn diagram showing the overlap of transcripts predicted to contain coding sequences by CPC2, CPAT, and CodAn. Numbers of total predictions and overlaps between programs are indicated.
**c**, Genomic location of novel transcripts relative to the GENCODE vM25 annotation, classified as classes 1–6. The schematic representation of each class is shown on the right. Class 1, novel transcript located within an intron of an annotated gene; class 2, novel transcript likely resulting from polymerase overrun; class 3, novel transcript sharing an intron with an annotated gene on the opposite strand; class 4, novel and annotated transcripts with overlapping exons on opposite strands; class 5, annotated gene located within the boundary of the novel transcript; class 6, novel transcript located in the intergenic region.
**d**, **e**, Sequence alignments of predicted protein products from novel transcripts MSTRG.1119.1 and MSTRG.10780.1 showing partial homology to arf-GAP and HGH1 protein sequences, respectively. The color key reflects the average BLOSUM62-based similarity of each residue to all others at the same alignment position: blue indicates higher similarity, green indicates lower similarity.

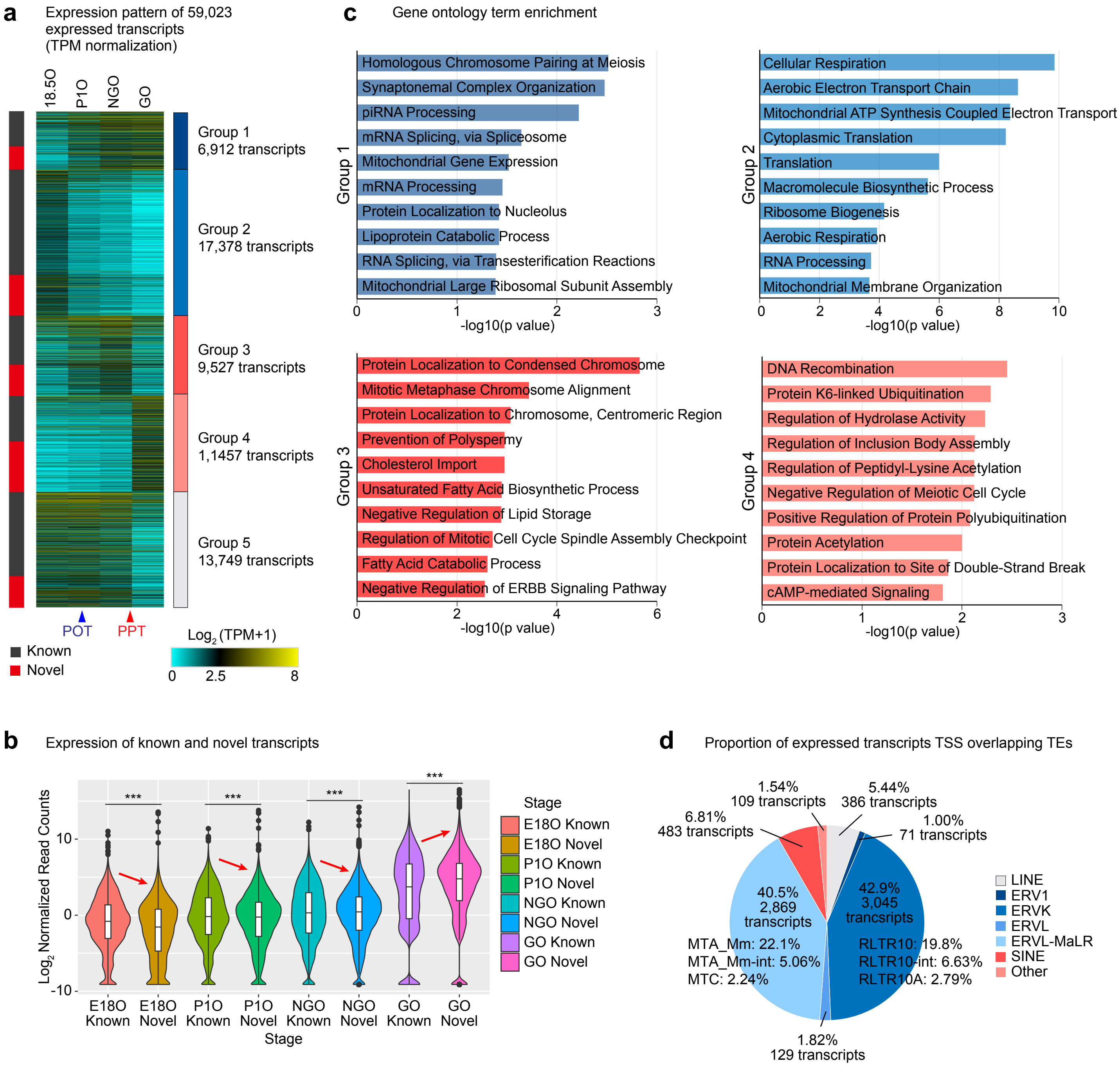


**Extended Data Figure 3. Expression profiles of novel transcripts.**

**a**, Heatmaps showing transcript expression during perinatal oogenesis (>1 TPM in at least one of the four developmental stages). Expressed transcripts were grouped into five k-means clusters.

**b**, Violin plots with a box plot showing log2-transformed transcript expression normalized by ERCC spike-in during perinatal oogenesis. Central lines represent medians; boxes indicate the interquartile range (IQR; 25th–75th percentile); whiskers extend to 1.5× IQR from the hinges. ***$P < 0.001$, Wilcoxon rank-sum test.

**c**, Gene ontology term enrichment analysis of genes in each group shown in Figure 1e.

**d**, Pie charts showing the proportion of TE families and types among TEs overlapping with TSSs of expressed transcripts.

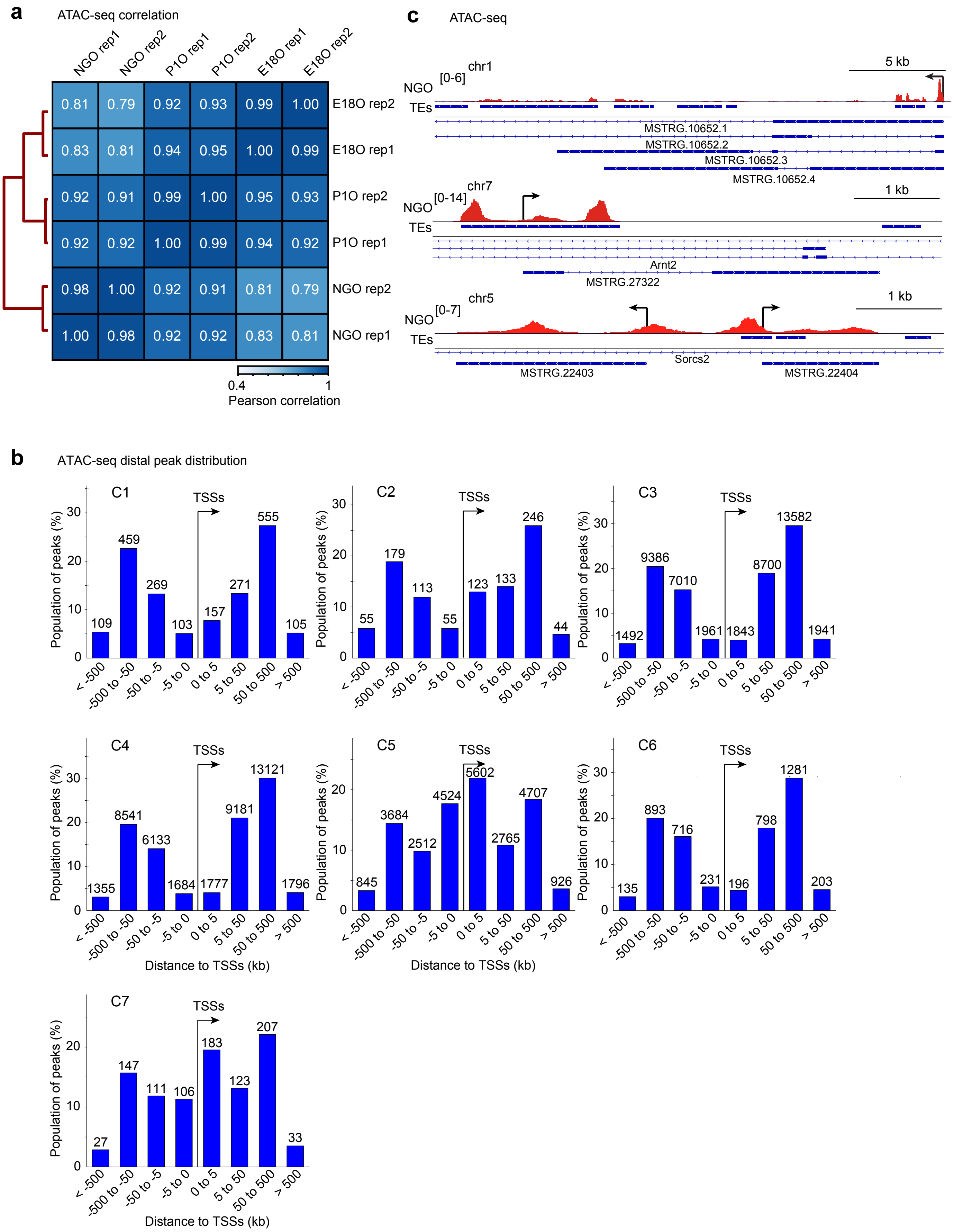
a
ATAC-seq correlation
NGO rep1
NGO rep2
P1O rep1
P1O rep2
E18O rep1
E18O rep2
0.81 0.79 0.92 0.93 0.99 1.00 E18O rep2
0.83 0.81 0.94 0.95 1.00 0.99 E18O rep1
0.92 0.91 0.99 1.00 0.95 0.93 P1O rep2
0.92 0.92 1.00 0.99 0.94 0.92 P1O rep1
0.98 1.00 0.92 0.91 0.81 0.79 NGO rep2
1.00 0.98 0.92 0.92 0.83 0.81 NGO rep1
0.4
1
Pearson correlation
c
ATAC-seq
chr1
[0-6]
NGO
5 kb
TEs
MSTRG.10652.1
MSTRG.10652.2
MSTRG.10652.3
MSTRG.10652.4
chr7
NGO
[0-14]
1 kb
TEs
Arnt2
MSTRG.27322
chr5
NGO
[0-7]
1 kb
TEs
Sorcs2
MSTRG.22403
MSTRG.22404
b
ATAC-seq distal peak distribution
C1
TSSs
Population of peaks (%)
109 459 269 103 157 271 555 105
C2
TSSs
55 179 113 55 123 133 246 44
C3
TSSs
1492 9386 7010 1961 1843 8700 13582 1941
C4
TSSs
1355 8541 6133 1684 1777 9181 13121 1796
C5
TSSs
845 3684 2512 4524 5602 2765 4707 926
C6
TSSs
135 893 716 231 196 798 1281 203
C7
TSSs
27 147 111 106 183 123 207 33
< -500
-500 to -50
-50 to -5
-5 to 0
0 to 5
5 to 50
50 to 500
> 500
Distance to TSSs (kb)

**Extended Data Figure 4. ATAC-seq analysis during POT.**
**a**, Heatmaps showing Pearson correlations of read counts among biological replicates in ATAC-seq datasets.
**b**, Bar chart showing the genomic distribution of ATAC-seq peaks according to their distance from TSSs.
**c**, Representative track views showing ATAC-seq signals at gene loci with accessible TEs downstream of TSSs. Arrows indicate the direction of transcription.

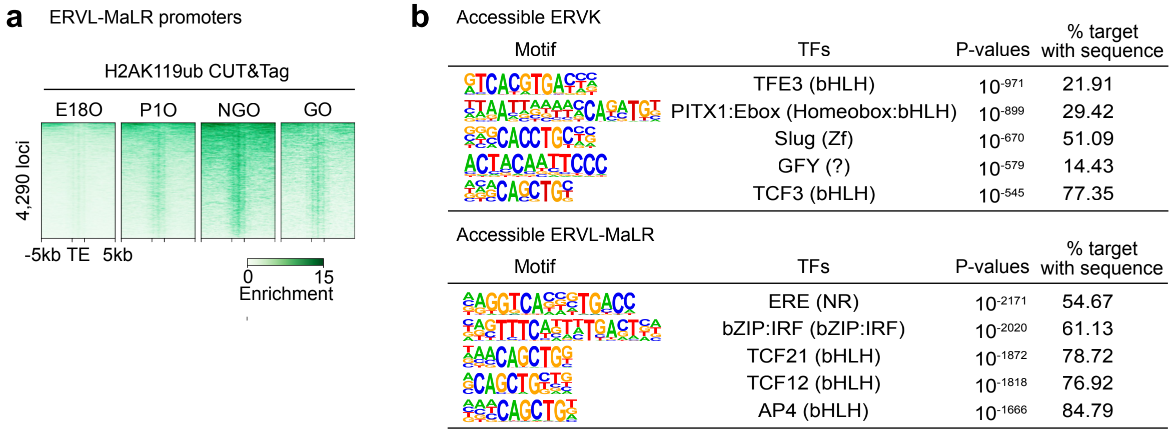


**Extended Data Figure 5. Analysis of ERVs in accessible chromatin regions.**

**a**, Heatmaps showing H2AK119ub enrichment at proximal accessible ERVL-MaLR regions during perinatal oogenesis.
**b**, HOMER known motif analyses of accessible ERVK (top) and ERVL-MaLR (bottom) loci in NGO.

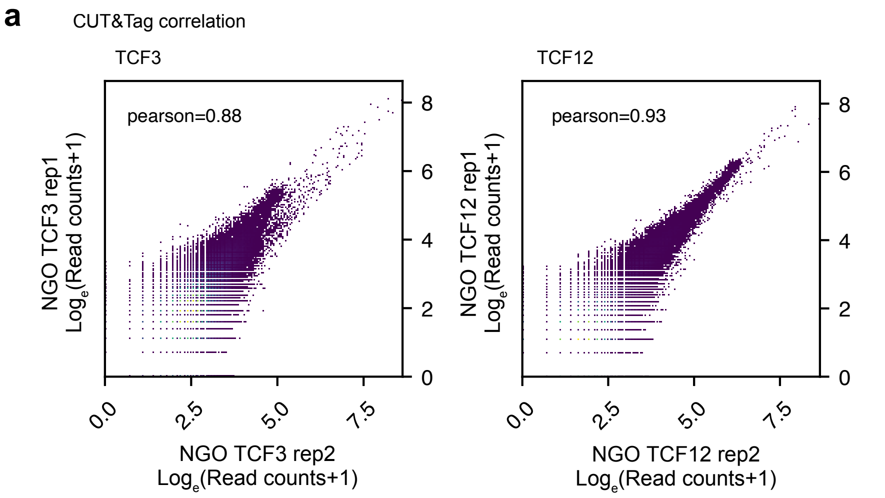


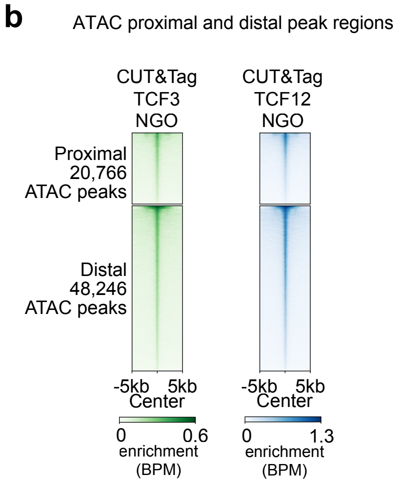


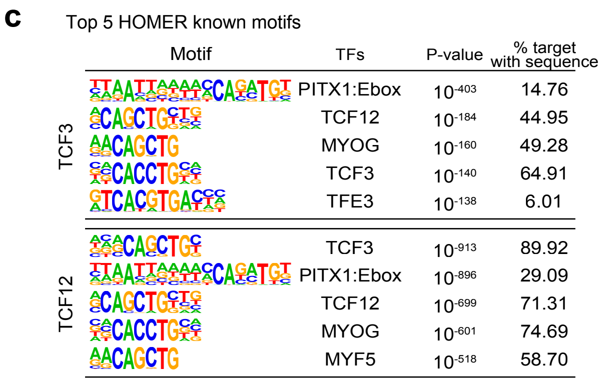


**Extended Data Figure 6. TCF3 and TCF12 CUT&Tag analysis in NGO.**

**a**, Scatter plots showing reproducibility between biological replicates in TCF3 (left) and TCF12 (right) CUT&Tag datasets. Pearson correlation coefficients are indicated in the panels.

**b**, Heatmaps showing TCF3 (left) and TCF12 (right) CUT&Tag enrichment dynamics at proximal (TSS ±1 kb) and distal (>1 kb from TSSs) ATAC-seq peaks shown in Figure 2c.

**c**, HOMER known motif analyses of TCF3 (top) and TCF12 (bottom) CUT&Tag peaks.

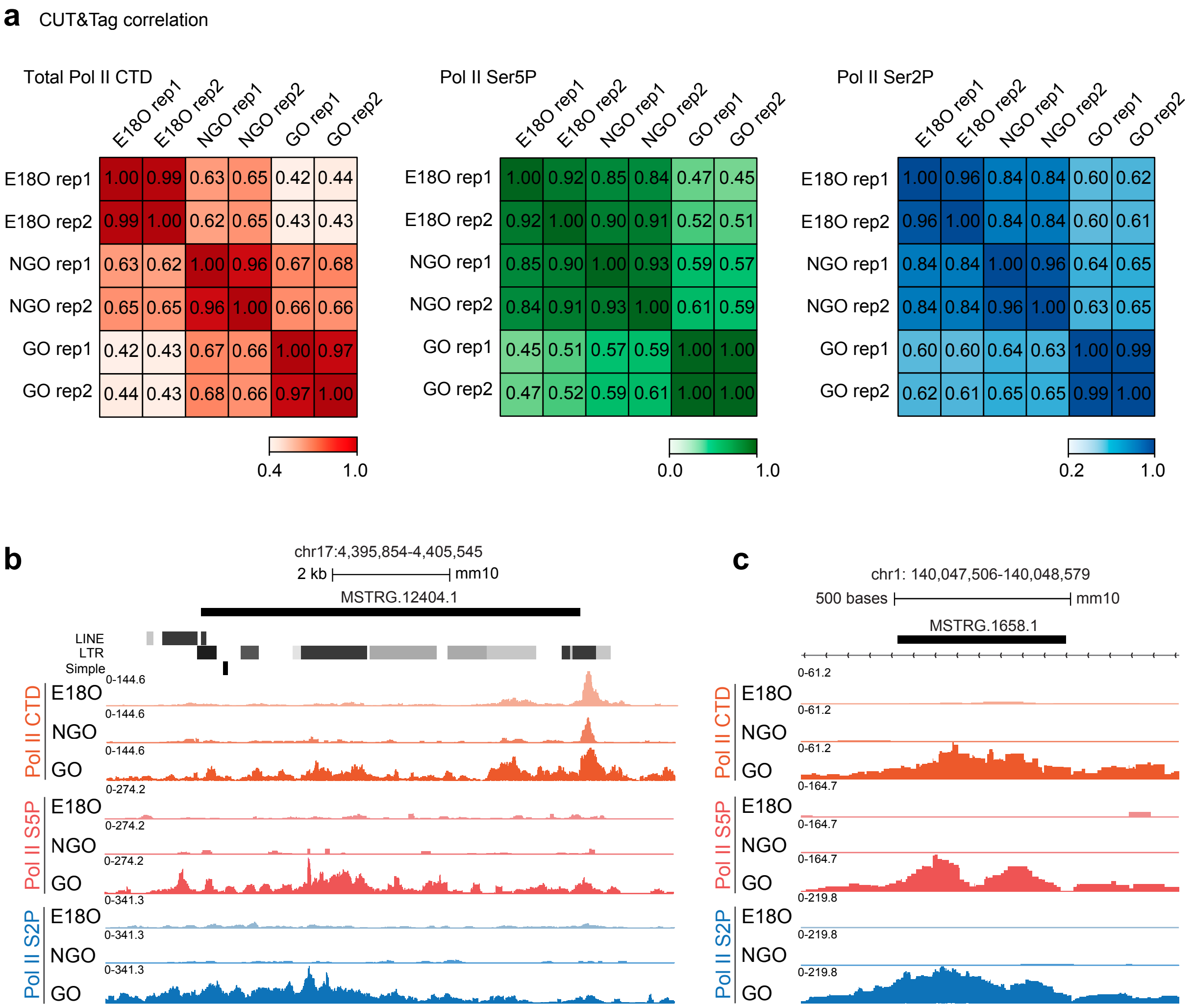


**Extended Data Figure 7. Pol II CUT&Tag analysis during perinatal oogenesis.**

**a**, Heatmaps showing Pearson correlations of read counts among biological replicates in total Pol II (left), Pol II Ser5P (center), and Pol II Ser2P (right) CUT&Tag datasets. Pearson correlation coefficients are indicated in the panels.

**b**, **c**, Track views showing total Pol II, Pol II Ser5P, and Pol II Ser2P enrichment at mono-exonic transcripts (MSTRG.12404.1 in **b** and MSTRG.1658.1 in **c**) derived from novel genes identified in this study during perinatal oogenesis.
